# Peptide Mold: A Novel Strategy for Mapping Potential Binding Sites in Protein Targets

**DOI:** 10.1101/2024.02.28.582665

**Authors:** Pritam V. Bagwe, Yogesh Jagtap, Vaibhav Ghegade, Janvhi Machhar, Elvis Martis, Shreerang V. Joshi, Prashant S. Kharkar

## Abstract

A novel concept titled ‘Peptide Mold’ for mapping potential binding sites in protein targets is presented. A large multiconformer tetrapeptide library comprising of 32 million conformations of all possible combinations of naturally-occurring amino acids was constructed and used for molecular docking analysis in the substrate-binding site of SARS-CoV-2 PLpro enzyme. The top-ranking, structurally-diverse tetrapeptide docked conformations (symbolizing peptide mold, analogous to a clay mold) were used then for elucidating a five-point pharmacophore. Ligand-based virtual screening of a large, multiconformer library of phytoconstituents using the derived five-point pharmacophore led to identification of potential binders for SARS-CoV-2 PLpro at its substrate-binding site. The approach is based on generating the imprint of a macromolecular binding site (cavity) using tetrapeptides (clay), thereby generating a reverse mold (with definitive shape and size), which can further be used for identifying small-molecule ligands matching the captured features of the target binding site. The approach is based on the fact that the individual amino acids in the tetrapeptide represent all possible drug-receptor interaction features (electrostatic, H-bonding, van der Waals, dispersion and hydrophobic among others). The ‘peptide mold’ approach can be extended to any protein target for mapping the binding site(s), and further use of the generated pharmacophore model for virtual screening of potential binders. The peptide mold approach is a robust, hybrid computational screening strategy, overcoming the present limitations of structure-based methods, e.g., molecular docking and the ligand-based methods such as pharmacophore search. Exploration of the peptide mold strategy is expected to yield high-quality, reliable and interesting virtual hits in the computational screening campaigns during the hit and lead identification stages.

## Introduction

Severe acute respiratory syndrome coronavirus 2 (SARS-CoV-2) is a novel coronavirus that emerged in Wuhan, China in December 2019 and has since spread globally. As of August 14, 2023, there have been 769,369,823 confirmed cases of COVID-19 worldwide and 6,954,336 deaths attributed to the disease, according to data from the World Health Organization (WHO).^1^ Since the beginning of the pandemic, various drugs have been used to treat COVID-19 patients, but no specific therapeutic drugs have yet been identified.^2^ The efficacy of drugs like remdesivir, chloroquine and hydroxychloroquine, ribavirin, ritonavir-lopinavir, favipiravir, oseltamivir, umifenovir, azithromycin, tocilizumab, and dexamethasone continue to be controversial.^3^ However, researchers kept on exploring repurposing and optimizing existing drugs for COVID-19 treatment, along with new drug discovery and development strategies.^4–6^ In addition to drug therapies, vaccines have been developed and deployed in various parts of the world. As of August 14, 2023, 50 vaccines against SARS-CoV-2 have been approved, and 242 vaccine candidates are under development with a total of 821 ongoing vaccine trials globally in progress.^7^ The Pfizer-BioNTech (BNT162b2)^8^ vaccine, Moderna (mRNA-1273)^9^ vaccine, Janssen (Ad26.COV2.S)^10^ vaccine, Oxford/AstraZeneca (Vaxzevria)^11^ vaccine, and the Sinovac (CoronaVac)^12^ vaccine are among the most widely used vaccines.

Despite the availability of vaccines, there are several factors that make it unlikely that vaccination alone will be sufficient to tackle the COVID-19 and any future pandemics. Firstly, vaccines are not 100% effective, and breakthrough infections continue to occur. Secondly, it is not yet clear whether vaccinated individuals can transmit the virus to others. Thirdly, there is a significant population worldwide that remains unvaccinated, and it will take time to achieve herd immunity through vaccination. In addition, new variants of SARS-CoV-2 keep emerging periodically. Understanding the molecular characteristics of these variants is central to finding a long-standing solution to the present problem.^13^ Researchers continue to study the efficacy of COVID-19 vaccines against the virus and its variants. Recent studies have shown that vaccination can significantly reduce the risk of severe disease, hospitalization, and death. However, there is ongoing research to understand how long the protection from vaccines lasts and whether booster shots will be needed.^14^ Other issues include long COVID, pediatric COVID, transmission of disease and many others. Overall, SARS-CoV-2 continues to be a significant public health threat, and researchers are working tirelessly to identify safe and effective therapies to control the pandemic.^15^ Extensive and continued research efforts are needed for finding efficacious treatments for COVID-19 and emerging viral infections.

Various targets for anti-SARS-CoV-2 drug development have been identified, including papain-like protease (PLpro), with cysteine protease and deubiquitinase catalytic activities, which is encoded inside the largest multi-domain non-structural protein 3 (Nsp3) of the SARS-CoV-2 genome.^16^ The PLpro enzyme is involved in all stages of the viral life cycle and plays a critical role in inhibiting the host immune response.^17–20^ It catalyzes the cleavage of the viral polyproteins, a crucial step in the replication and transcription of the virus. Therefore, it is proposed that inhibition of PLpro enzyme could potentially reduce the viral load in infected cells and prevent the virus from spreading. In fact, it has been demonstrated that PLpro inhibition not only stopped the viral replication, but also significantly boosted the host immune response, making it a promising target for drug discovery and development.^21^ Significant number of drugs have been studied for their potential to inhibit PLpro, including HIV protease inhibitors, repurposed medications like sitagliptin and daclatasvi, and investigational drugs like tropifexor, ebselen, disulfiram, tanshinone, cryptotanshinone, asunaprevir, simeprevir, famotidine, carmofur, PX12, and tideglusib.^22–24^ Several new chemical entities (NCEs) belonging to varied chemical series have been reported as potent or moderately potent inhibitors of PLpro^25,26^ (reviewed in Ref. 25). Few of these NCEs such as GRL067, VIR250, VIR251, XR8-24, and XR8-23, are currently under detailed biological testing or clinical investigations.^25^ Combining GRL0617 with a sulfonium-tethered peptide generated from the PLpro-specific substrate LRGG led to a novel peptide-drug combination.^27^ Phytochemicals such as fatty acids and natural biflavones have been confirmed to be effective PLpro inhibitors.^28^ Natural products such as hypericin, rutin, and cyanidin-3-*O*-glucoside have also been identified as PLpro inhibitors by a combination of in silico studies and in vitro enzyme inhibition assays.^29^ However, further research is needed to determine their efficacy and safety for treating COVID-19.

The PLpro enzyme of SARS-CoV-2 has been extensively studied from the structural bioinformatics and molecular design perspectives.^30,31^ More than 50 crystal structures (including mutant enzyme) have been reported till date. The PLpro structure constitutes six domains, namely, finger subdomain, blocking loop, catalytic triad, palm, thumb, and ubiquitin-like subdomain.^32,33^ Several crucial residues, including Cys111, His272, Asp286, and Tyr268, are involved in the overall function and structural integrity of the enzyme. The zinc-binding domain is also essential for maintaining its structural integrity. Effective inhibitors need to target the hydrophobic active site of PLpro, which includes Asp164, Arg166, Met208, Pro247, Pro248, Tyr264, Tyr268, Gln269, Tyr273, and Thr301. Inhibitors must also suppress or inhibit the proteolytic, deubiquitinating, and deISGylating activities of the coronavirus PLpro protease. These crystal structures of PLpro are extremely helpful for the structure-based design of novel inhibitors.

Conventionally, protease inhibitors are either peptides or peptidomimetics derived from the peptide substrate(s). The features of the substrate peptides around the cleavage sites are imprinted along with binding-affinity enhancing groups in the potential inhibitor structures. The availability of a large number of rationally-designed peptide inhibitor drugs, particularly antiviral drugs such as HIV protease inhibitors, clearly endorsed the design approach based on mimicking the interactions within the substrate binding site.^34^ These sites are best tuned to recognize and bind peptide substrates with unique amino acid sequences, making them highly selective/specific. Building on these molecular recognition elements, here we introduce a novel concept titled ‘peptide mold’, analogous to epitope mapping in immunology. The core of the workflow involved the virtual screening of all possible tetrapeptides (20^4^ = 160,000) derived from naturally occurring L-amino acids, for potential binding within the protease substrate-binding site. One of these tetrapeptides would exactly match the substrate sequence around the cleavage site. The idea was to identify the tetrapeptides (treated as clay, which could take shape of the cavity due to conformational flexibility) binding more efficiently with respect to shape and electrostatic complementarity, within the substrate-binding site (treated as mold) using molecular docking, extract their pharmacophoric features from their binding conformations (clay shape) and use them for pharmacophore elucidation, followed by pharmacophore-based virtual screening of small-molecule databases for virtual hits identification. In other words, we attempted to identify tetrapeptide conformations, matching with the shape of substrate-binding site, but at the same time bind tightly due to their complimentary structural features contributing to the magnitude of various ligand-receptor interactions. The process tried to imprint the complimentary shape- and electrostatic features of the binding site simultaneously, giving the base shape, which could, in turn, be used for identifying potential binders matching with these complimentary imprinted substrate-binding site features, leading to a complimentary shape which could be used endlessly for virtual screening for potential binders within the substrate-binding site of the protease.

In the present work, we have applied a novel peptide mold concept to imprint the complimentary shape- and electrostatic features of the protease substrate-binding site and use them for virtual screening for potential binders of the target protease, i.e., SARS-CoV-2 PLpro enzyme. The top-ranking, structurally diverse, tetrapeptides thus obtained using molecular docking, were used for pharmacophore elucidation. The developed pharmacophore was then used for ligand-based virtual screening of a curated phytochemical library for identifying potential virtual hits, which could then be used for in vitro PLpro enzyme inhibition assays. The virtual screening results from direct docking the phytochemical library were compared with the peptide mold-based virtual screening. The process flow is presented in Figure 1. The concept is presented here taking PLpro as a test case, but can be extended to any protein target since the target binding sites would best identify the complimentary peptidic ligands, e.g., tetrapeptides. The process can be used for ligand-based virtual screening of huge databases and can save computational resources and the ensuing cost which would otherwise be incurred during the structure-based virtual screening using molecular docking at multiple levels such as initial high-throughput, followed by standard and in the end, precise modes.

## Materials and Methods

Hardware and Software: All the molecular modeling studies described herein were performed on HP Laptop (Intel® Core™ i7-5500U CPU @ 2.40 GHz, RAM 4 GB) running Windows 8.1 Home Basic Operating System. Schrödinger Small-Molecule Drug Discovery Suite Release 2020-3 and products included therein (Maestro v12.5, LigPrep v2.2, Epik v5.3, Glide v8.8, Prime v4.3, Phase v3.5) were used for performing various molecular modeling operations.^35^ OpenEye Scientific Software Applications 2022 and the modules therein (OMEGA 4.2.1.1^36^, MakeReceptor 4.2.0.1^37^, FRED 4.2.0.1^38^) were used for conformer generation, minimization and docking. Few auxiliary softwares and tools were used and are mentioned in the respective sections.

### Tetrapeptides Multiconformer Library Generation

A library of tetrapeptides comprising 160,000 members (20 X 20 X 20 X 20), using naturally-occurring L-amino acids) was built from their SMILES notations using Maestro v12.5, followed by Ligand Preparation using LigPrep 2.2 (default settings, pH 7.4). The stereochemistry at αC was fixed to *S*. The prepared structures were imported in OMEGA 4.2.1.1 wherein further molecular modeling operations such as tautomer generation, partial charge calculation (AM1Bcc charges), and conformer generation (total of 200 each) and energy minimization were carried out. A library of 32,000,000 tetrapeptides conformers was then used for molecular docking step using FRED 4.2.0.1. Chart 1 depicts the overall process workflow.

### Molecular Docking of Tetrapeptides Multiconformer Library

SARS-CoV-2 PLpro crystal structure (PDB ID 6WX4^33^) was downloaded from Protein Data Bank.^39^ The peptide ligand was covalently bound Cys111 of PLpro. The covalent bond was broken and the corresponding Hs were added before executing the protein preparation task workflow using MakeReceptor 4.2.0.1 (default settings). The generated OEDesign Unit was used for molecular docking using FRED 4.2.0.1 (Scoring function: Chemgauss 4). The tetrapeptides multiconformer library containing 32 million members was docked in PLpro active site. Following the docking operation, top-scoring binding poses (total 500) were extracted and used for docking studies using ExtraPrecision (XP) mode in Glide v8.8.

**Chart 1.**
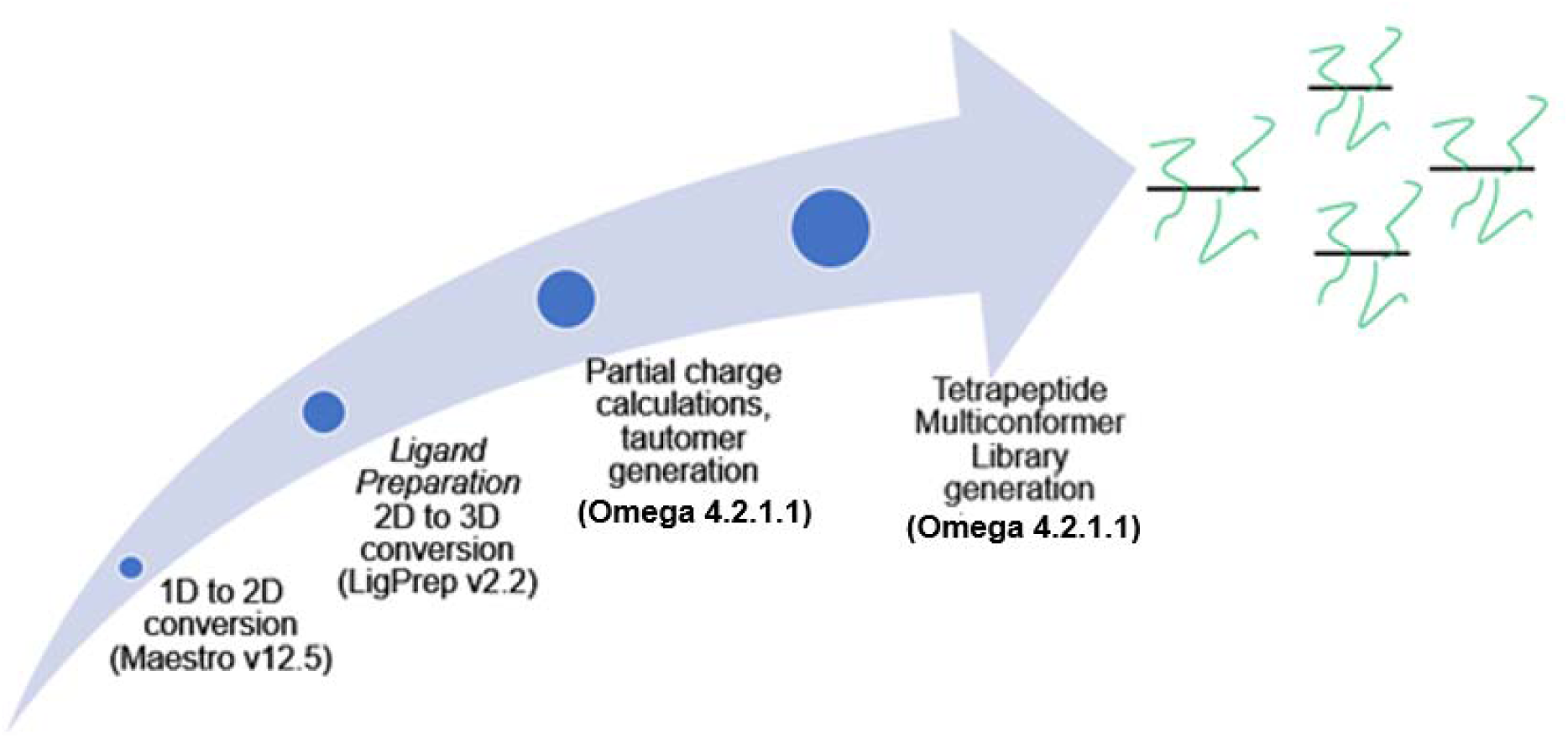
Steps involved in the generation of multiconformer library of tetrapeptides

### Molecular Docking of Top-Scoring Poses using Glide XP

The PLpro structure (PDB ID 6WX4) was imported in Maestro v12.5. The covalent bond of peptide ligand with Cys111 was broken and the corresponding Hs were added. The protein structure was then submitted to Protein Preparation wizard workflow (default settings). Missing side chains and loops, if any, were filled using Prime v4.3. Once completed, the problems were viewed and corrected, if necessary. For alternate positions, the position with higher occupancy (i.e., higher number) was committed. Other problems such as overlapping atoms, etc., are resolved in later steps. Nonligand Het atoms such as DMS, or other crystal structure adjuncts are deleted. Lone waters are selected and deleted. The modified protein was then further refined using PROPKA at pH 7.4 and optimized. Waters with less than two H-bond to non-waters were removed. Later, a restrained minimization was carried out using default settings (OPLS3 force field). The prepared protein was further subjected to Receptor Grid Generation (Glide v8.8) and the grid was then used for XP docking operation with the top-500 docked poses from FRED docking run in the active site of PLpro. All the default settings were used. The docked poses were minimized and RMSD to input ligand geometries were calculated. The top-100 docked poses were used for selection of top-10 most diverse tetrapeptides.

### Pharmacophore Generation from Top-10 Diverse Tetrapeptides

The top-10 most diverse tetrapeptides were used for pharmacophore generation. The diversity analysis was performed using MACCS Fingerprints as implemented in Molecular Operating Environment (MOE) 2022.^40^ The pharmacophore was generated using Phase v3.5. There are six in-built pharmacophoric features in Phase, namely H-bond acceptor (A), H-bond donor (D), hydrophobic (H), negatively-ionizable (N), positively-ionizable (P), and aromatic ring (R). The pharmacophore model was developed using a set of pharmacophore features to generate sites for all the 10 tetrapeptides. The alignment was measured using a survival score and the default values were used for the hypothesis generation. The pharmacophore hypothesis was selected based on the best fit value. The selected 5-point pharmacophore had all the necessary features such as H-bond donor as well as H-bond acceptor in addition to others.

### Phytochemicals Multiconformer Library Generation

For the pharmacophore-based screening, a large library of phytochemicals containing 158,920 members was used. The SMILES notations from the Excel file were imported in Maestro v12.5. The phytochemicals with molecular weight >700 Da were removed to save time during further molecular modeling operations such as conformer generation. The curated library of 148,324 phytochemicals was processed in the same way as tetrapeptide multiconformer library generation using LigPrep v2.2 and OMEGA 4.2.1.1 to generate 14,618,712 conformers.

### Pharmacophore-based Screening of Multiconformer Phytochemicals Library using 5-Point Pharmacophore Hypothesis and Molecular Docking of Top-five Hits

The multiconformer phytochemicals library generated using OMEGA 4.2.1.1 was screened using Phase pharmacophore hypothesis with default settings. Partial matching with pharmacophoric features (3/5) was permitted. The ranking of the hits was based on a number of features matched and the corresponding scores. The top-five hits were then subjected to molecular docking using MOE 2022.02 with default setting. Two crystal structures of SARS-CoV-2 PLpro (PDB IDs 6WX4 and 6WUU^33^) were used for docking analyses. The covalently-bound peptide ligands were treated as described previously, before the protein preparation step was executed. The lowest-energy conformations of the phytochemical virtual hits were then docked in PLpro catalytic site. The initial placement of the ligands (including crystal structure bound ligands) was done using the default Triangle Matcher method and initial scoring was done using London dG scoring methodology as implemented in MOE 2022.02. The post-placement refinement was done using induced fit strategy and the refined poses were finally scored with the help of GBVI/WSA dG scoring methodology. Top-five scoring poses for each of the five hits from pharmacophore search were further analyzed for interaction with catalytic site of PLpro. The hits were further subjected to molecular, physicochemical and ADME/T property evaluation using QikProp utility as implemented in Schrödinger Small-Molecule Drug Discovery Suite Release 2020-3.

### Molecular Dynamics (MD) Simulations of the Top-ranking Hits from Pharmacophore Screening of Multiconformer Phytochemicals Library

MD simulations can provide valuable insights into the binding kinetics, thermodynamics, and structural changes that occur during ligand-receptor interactions. To compute the binding affinity of the tetrapeptides to SARS-CoV-2 PLpro, we performed all-atom unbiased molecular dynamics simulations using the AMBER22 suite of programs which contain all the necessary tools for the generation of parameter-topology files and the analysis thereof.

**Figure 1.**
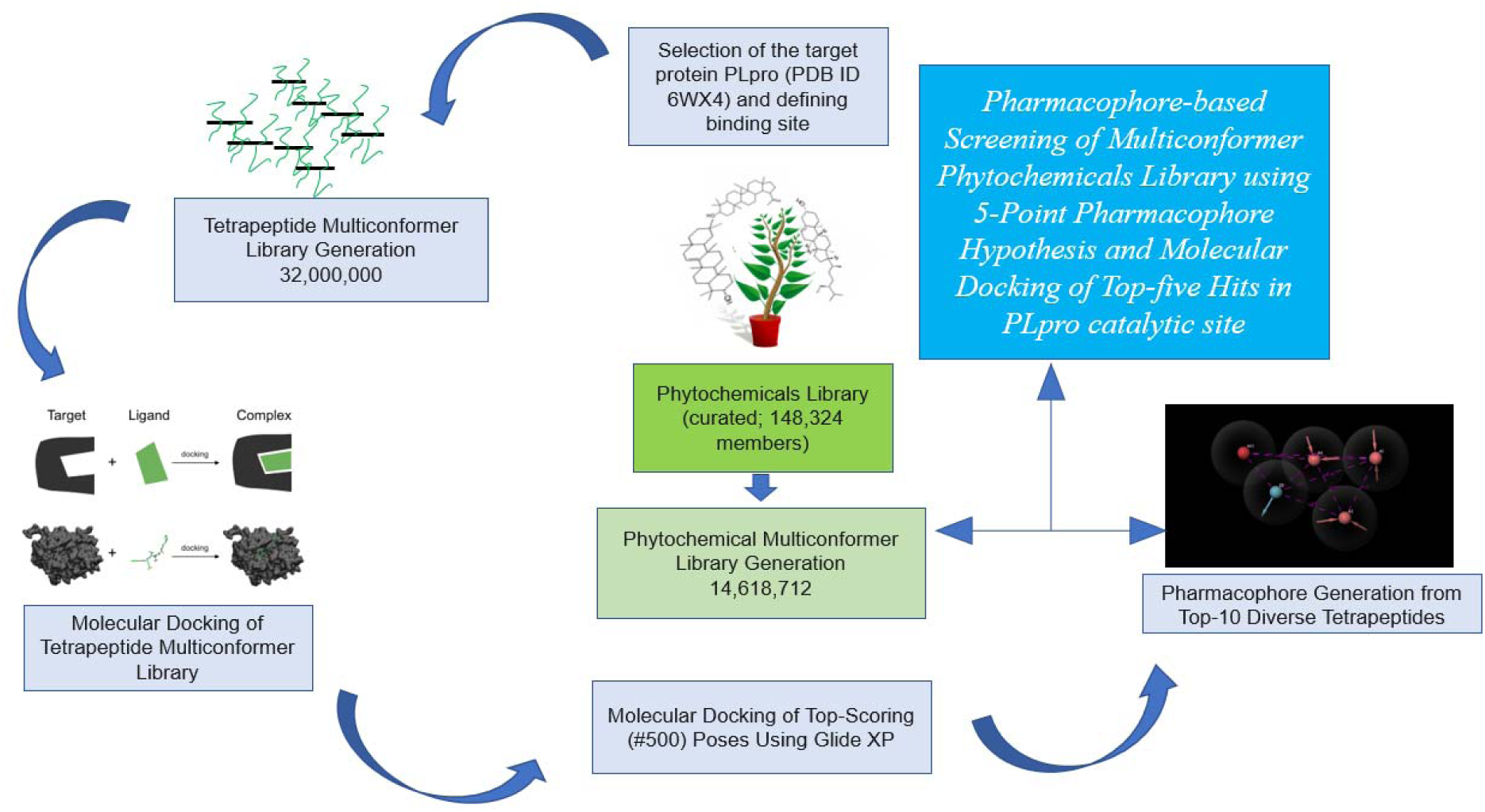
Process flow of the peptide mold concept

The initial coordinates for MD simulations were obtained from the molecular docking and scoring exercise. The system was prepared in the leap module in AMBERTools22. The protein atoms were assigned force field parameters from the ff14SB library and the parameters for the zinc coordinate bonds were obtained from the ZAFF force field library.^41^ Atom charges non-protein atoms were computed using the AM1-BCC^42,43^ method in the antechamber module in AMBERTools22. This module helps to define the atom type, bond type, and atomic equivalence, and to generate the initial coordinate file. Antechamber and parmchk generate an additional force field file that writes the missing parameters from GAFF2^44^, which is used as an input file for the leap. This reads in the force field, topology, and coordinate information and produces files necessary for further calculations which include the parameter-topology file. The TIP3P water model^45^ was used to solvate the complexes in a periodic truncated octahedron box; the edge of the solvent box was kept 10 Å away from any part of the complexes. The system was neutralized by adding counter ions Na+ or Cl-around the complex. The complexes were minimized using three cycles of minimization for a total of 50,000 steps. The first 5,000 steps were steepest descents, and the remaining 45,000 steps were conjugate gradients with the cutoff for non-bonded interactions placed at 9.0 Å.

The first cycle of minimization was performed with a 25 kcal/mol Å restraining force on the solute, this force was lowered to 5 kcal/mol Å in the next cycle of minimization. The final cycle of minimization was carried out without any restraining forces on the solute as well as the solvent. Following a thorough geometry optimization, the system was gradually heated from 0 K to 300 K for 1 ns using a 2 fs integration time-step under the NVT ensemble with a harmonic restraining force of 50.0 kcal/(mol LJ) placed on the solute atoms. The heating stage consisted of two phases; for the first 50 ps, the temperature was linearly raised to 300 K, and in the second phase, the temperature of the system was held steady at 300 K for the remaining time by Langevin dynamics and a collisional frequency of 2.0 ps-1.^46,47^ Further, the system was equilibrated under the NPT ensemble in two stages. In the first stage, the system was equilibrated for 1 ns using a 2 fs integration time-step with a harmonic restraining force of 5.0 kcal/(mol LJ) on the solute atoms. In the second stage, the system was allowed to equilibrate without any restraining force for a period of 1 ns using a 2 fs integration time-step. These fully equilibrated systems were simulated for 3 independent production cycles of 100 ns each, amounting to a total sampling time of 300 ns per system. The GPU-enabled version PMEMD^48–50^ was used for MD simulations running on an RTX3080 GPU card. For the production stage, Newton’s equation of motion was solved with an integration time step of 4 fs using the Hydrogen Mass Repartitioning scheme.^51^ The long-range electrostatic interactions were evaluated by the Particle Mesh Ewald method^52^ and all non-bonded interactions were truncated beyond 10 Å. All bonds involving the hydrogen atoms were restrained using the SHAKE algorithm.^53–55^

The simulations were carried out at 300 K and 1 bar. A total of 15,000 snapshots from each independent production cycle were saved for analysis. Analysis of the trajectories, the root-mean-squared deviation from the starting geometry, and the mean atomic fluctuations were calculated using the CPPTRAJ tools^56^ in AMBERTools22. The binding free affinity of the tetrapeptides to the SARS-CoV-2 PLpro was computed using the molecular mechanics Generalized Born surface area (MMGBSA) of each complex generated by the MD simulation.^57^ The solvent dielectric constant was at 80 and the solute dielectric constant was set to 1.0. The atomic radii for all the atoms were set to mbondi3. In MMGBSA calculation the gas-phase interaction energy (ΔEMM) and non-polar solvation energy (ΔGNP) are calculated. The electrostatic solvation energy (ΔGGB) was calculated using the GB model developed by Onufriev and co-workers^58^ (igb = 5 also known as the GBOBC model).

### Direct docking of Phytochemicals Library in PLpro Substrate-binding Site

Molecular docking studies of the phytochemical’s library were performed on Centre for Development of Advanced Computing (CDAC) (Pune, India) servers. Jobs were submitted on CDAC servers using – 1. Dell Laptop (Intel® Core™ i5-8250U CPU @ 1.60, 1.80 GHz, RAM 8 GB) running Windows 8.1, 64-bit Operating System, x64-based processor and radeon graphics, 2. Dell Laptop (Intel® Core™ i3-8145U CPU @ 2.10, 2.30 GHz, RAM 8 GB) running Windows 8.1, 64-bit Operating System, x64-based processor, and 3. Lenovo Laptop (Intel® Core™ i5-9300H CPU @ 2.40 GHz, RAM 8 GB) running Windows 8.1, 64-bit Operating System, x64-based processor.

Discovery Studio Visualizer v20.1.0.19295, 2019^59^ and the products included therein were used for performing various molecular modeling operations described below.

1. *Protein preparation.* The crystal structure of SARS-CoV-2 PLpro complexed with its inhibitor (PDB ID: 6WX4) was imported. CHARMm force field was selected for further modeling operations. Hydrogen atoms were added from the *Chemistry* tab. Later, the protein preparation protocol was executed at pH 7.4 comprising of tasks such as modeling loop regions, inserting missing atoms in incomplete residues, deleting alternate conformations, protonating titratable residues and removing water molecules and hetero atoms which might be trapped in the crystal structure during the crystallization procedure. Following the completion of protein preparation, the prepared protein was subjected to minimization using *Smart Minimizer* algorithm. On completion, the file is saved as a new entry in DDS format.
2. *Defining binding site*. The prepared protein file was opened and from the *Define Binding Site* tab, the binding site with bound ligand option was selected. A translucent sphere appeared on the ligand, the size of which was expanded twice. The native ligand was deleted and the coordinates of the defined binding site were noted for further processing.
3. *Phytochemicals ligand preparation.* Previously prepared Phytochemicals library was imported in the Discovery Studio interface and used for next set of studies.
4. *LibDock (HTVS) and result analyses*. The protein-ligand docking and visualization studies were performed using the LibDock program in Discovery Studio. LibDock^60^ is a high throughput algorithm for docking ligands into active receptor sites. Ligand conformations were aligned to polar and apolar receptor interaction sites (hotspots) and the best scoring poses were reported. Ligand conformations were pre-calculated during ligand preparation protocol. The prepared and minimized structure to be used as target receptor (with defined binding site) was selected as input receptor. The ligands to be docked were selected in the input ligands option. The coordinates obtained while defining the binding site sphere were mentioned in the input sphere option. Docking preferences were set to user specified. Max hits to save, max no. of hits, max conformations hits, max start conformations options were all set to 1, since conformations were already generated. Also, the conformation method was set to none. Rest other parameters were set to default and the protocol was run. Both the target and ligand are typed in CHARMm forcefield before initializing the docking protocol. After the run finished, analyses of the docking results were performed. LibDock score was obtained as a numerical value scoring function for each docked ligand.
5. *Calculating Binding Energy*. The binding free energy was calculated using CHARMm force field. The binding free energy was estimated between each ligand and the receptor. The free energy of binding for a receptor-ligand complex can be calculated from the free energies of the complex, the receptor, and the ligand. Using CHARMm based energies, it was possible to estimate these free energies and thus calculate an estimate for the overall binding free energy. The prepared and minimized structure to be used as target receptor was selected as input receptor. Top 8% ligands from the docking run were selected in the input ligands option. In-situ minimization was set to true and *Smart Minimizer* protocol was selected. The coordinates obtained while defining the binding site sphere were mentioned in the input sphere option. Partial charge estimation method was set to MMF and all other options were used as default. Parallel processing was set to true. The *Calculate Binding Energy* score is obtained as a numerical value scoring function for each docked ligand.

The top-5 hits obtained from direct docking of the phytochemicals library were subjected to molecular docking using MOE 2022.02 with default setting using the two crystal structures of SARS-CoV-2 PLpro (PDB IDs 6WX4 and 6WUU^33^) with same settings as described in earlier description for pharmacophore hits.

## Results and Discussion

The present study introduced a novel concept titled ‘peptide mold’ for hit discovery for a potential anti-SARS-CoV-2 target, PLpro enzyme. The study used biomimicry strategy based on tetrapeptides. The protease active site is tuned to identify specific amino acid sequences of the substrate protein for initial, highly-specific binding, followed by the precise catalytic events leading to selective peptide bond cleavage. We started off with a thought process that if we dock smaller peptides, e.g., tetrapeptides, in the PLpro active site, we would be able to identify potential binders based on their shape- and electrostatic complementarity with the active site residues. The tetrapeptides would contain the side chains of varied nature (polar/nonpolar, acidic/neutral/basic, hydrophilic/hydrophobic, aliphatic/aromatic, H-bond Don/H-bond Acc) which could also take up varied shapes due to the conformational freedom (dihedral angles θ and ψ). The thermodynamically-allowed conformational freedom and the nature of side chains of the constituent amino acids would make them ideal candidates for exploring the complimentary enthalpic and entropic contributions to the overall binding energy via ligand-receptor interaction forces, such as electrostatic, H-bonding, van der Waals, hydrophobic, cation-π, π-π stacking and few others. We attempted to mimic nature’s minimalistic design approach, i.e., we tried to imbibe biomimicry in our molecular design strategy.

Initial thought process was based on three options for smaller peptides, namely, tripeptides, tetrapeptides and pentapeptides, keeping in mind the overall size (molecular weight) of the resulting peptides. The molecular weights of the smallest tripeptide (Gly-Gly-Gly), tetrapeptide (Gly-Gly-Gly-Gly) and the pentapeptide (Gly-Gly-Gly-Gly-Gly) are 189.17, 246.22 and 303.27 Da, respectively. The corresponding values for the largest tripeptide (Trp-Trp-Trp), tetrapeptide (Trp-Trp-Trp-Trp) and pentapeptide (Trp-Trp-Trp-Trp-Trp) are 576.65, 762.87 and 949.08 Da, respectively. For a drug-like chemical space, the tetrapeptides were obviously the best option. Another reason to choose a tetrapeptide library was due to the preference of PLpro for a specific tetrapeptide sequence, Ac-LRGG-ACC, for recognition and cleavage; the same sequence is also recognized by other deubiquitinating enzymes. We, therefore, initiated the generation of tetrapeptide library consisting of all possible combinations of naturally-occurring L-amino acids (absolute stereochemistry *S*) (Chart 1).

The processed tetrapeptide library of 32 million conformations derived from 160,000 tetrapeptides was docked into the active site of PLpro. The native ligand was covalently bound to the active site nucleophilic Cys111 (cleavage site). For the docking tool to identify Het groups, thereby identifying the site for docking, the covalent bond with Cys111 was broken and the corresponding Hs added. The generated receptor unit was used for docking of the multiconformer tetrapeptide library. The task was broken into smaller subjobs, consisting of 8000 tetrapeptides (160,000 conformations); each docking run generated top-500 hits (including multiple conformations of same molecule). These 10,000 (500X20) docked poses were further docked using same workflow to select the top-500 poses. Table 1 lists the top-10 tetrapeptides with their corresponding FRED Chemguass4 scores.

As seen from Table 1, the top-10 tetrapeptides exhibited unique structural features such as – 1. Presence of Gly as the C-terminus residue, 2. Presence of Gly as the second residue in 9/10 cases, 3. presence of Ala/Ser as the N-terminus residue in 8/10 cases, and 4. presence a hydrophobic residue such as Phe (4/10), Leu (2/10), Thr (2/10) and Tyr (2/10 cases).

Interestingly, the recognition sequence specific to PLpro cleavage site features were retained by the top-scoring tetrapeptides. The average relative size (mol wt) of the top-scoring tetrapeptides was 364.18, although bit skewed by peptide No. 9 (Table 1). The peptide mold workflow, thus, could successfully identify a unique set of tetrapeptides in the top-scoring poses.

**Table 1.**
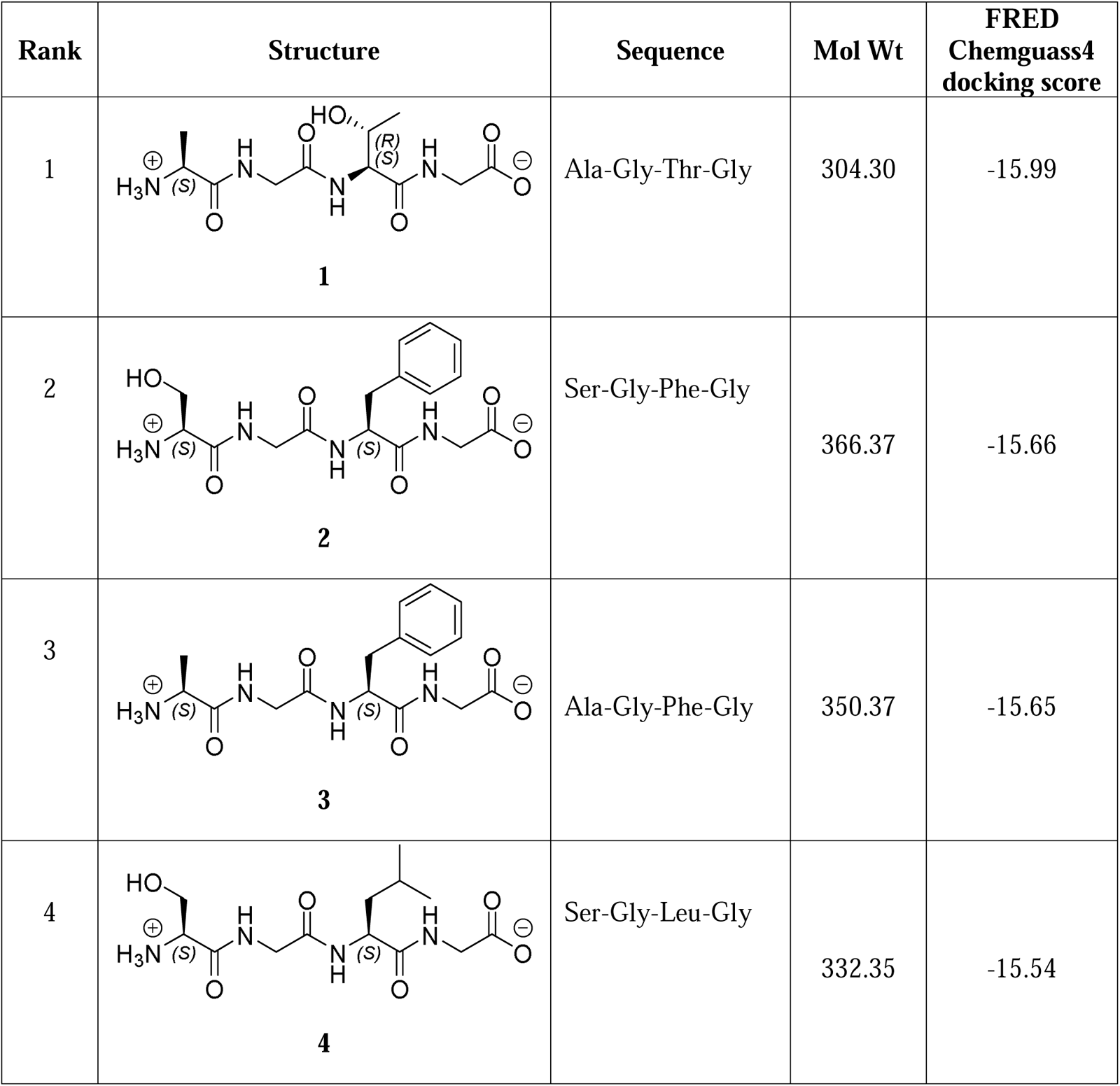
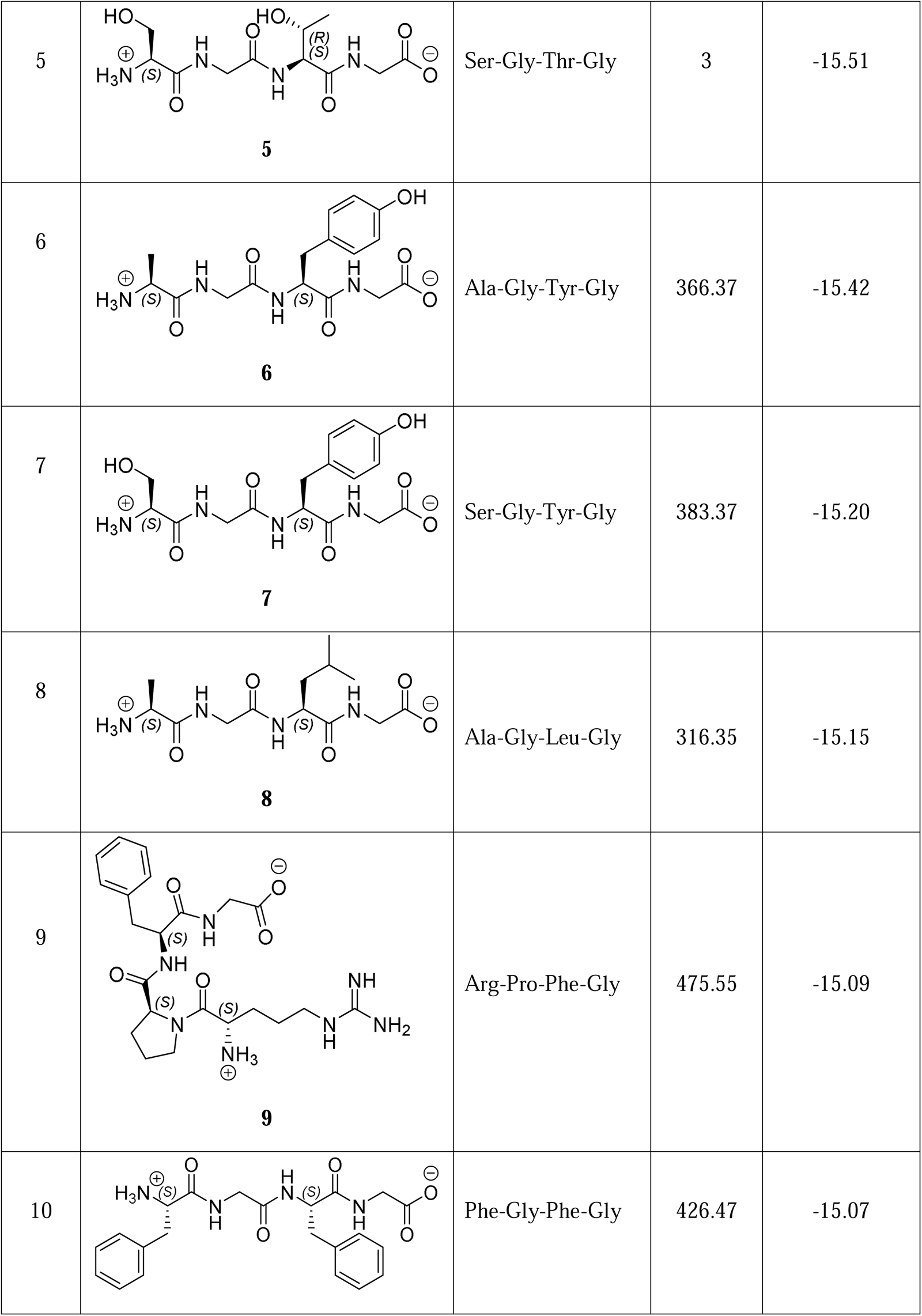
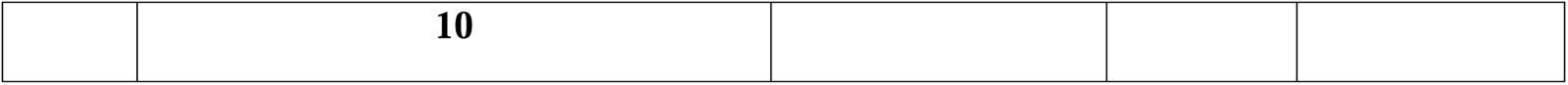
Representative top-scoring tetrapeptides with their molecular properties and docking scores.

| Rank | Structure | Sequence | Mol Wt | FRED<br>Chemguass4<br>docking score |
| --- | --- | --- | --- | --- |
| 1 | **1** | Ala-Gly-Thr-Gly | 304.30 | -15.99 |
| 2 | **2** | Ser-Gly-Phe-Gly | 366.37 | -15.66 |
| 3 | **3** | Ala-Gly-Phe-Gly | 350.37 | -15.65 |
| 4 | **4** | Ser-Gly-Leu-Gly | 332.35 | -15.54 |
| 5 | **5** | Ser-Gly-Thr-Gly | 3 | -15.51 |
| 6 | **6** | Ala-Gly-Tyr-Gly | 366.37 | -15.42 |
| 7 | **7** | Ser-Gly-Tyr-Gly | 383.37 | -15.20 |
| 8 | **8** | Ala-Gly-Leu-Gly | 316.35 | -15.15 |
| 9 | **9** | Arg-Pro-Phe-Gly | 475.55 | -15.09 |
| 10 |  | Phe-Gly-Phe-Gly | 426.47 | -15.07 |
|  | **10** |  |  |  |

The docked pose of top-scoring tetrapeptide **1** (Table 1) in the active site of PLpro is shown for reference (Figure 2). The active site shape was adapted by **1**, which interacted with PLpro via four H-bonds with Gly163 (two H-bonds with backbone –NH and –C=O groups), Gly271 and Tyr268 (one H-bond each with backbone –NH and –C=O groups, respectively). Additionally, the N-terminus –NH_3_ of tetrapeptide formed a strong salt-bridge interaction with anionic side chain of Asp164. The 3^rd^ residue of **1** was exposed to solvent. The crystal structure ligand and **1** overlapped in the PLpro active site in similar fashion (pose not shown). The C-terminal Gly of **1** was located in proximity to the catalytic Cys111, confirming the appropriate occupancy of the catalytic site of PLpro.

The top-500 docked tetrapeptides were further docked in the PLpro active site using Glide Extra Precision (XP) mode. The top-scoring pose of each of the tetrapeptides was sorted as per Glide XP score. The top-100 docked poses were subjected to diversity analysis based on MACCS fingerprints. The top-10 most diverse tetrapeptides were then taken up for pharmacophore elucidation using Phase v3.5. A five-point pharmacophore comprising of AAADN features, was generated using default settings. Figure 3 depicts these pharmacophoric features and their geometric arrangement, including directions of the A and D features. The top-five hits from Phase pharmacophore screening were mapped onto the five-point pharmacophore and are shown in Figures 4a-e. Other related pharmacophore hypotheses were not discussed here due to closeness (in terms of number of features and their nature) with the top-scoring hypothesis. The selected pharmacophore hypothesis (AAADN) was used for virtual screening of the multiconformer phytochemicals library using Phase v3.5 (default settings). In summary, the active site of PLpro was successfully mapped in the form of 5-point pharmacophore, ready to be deployed for the hit identification in the ensuing virtual screening exercise.

The molecular structures of top-five hits, **11**-**15** from Phase virtual screening of multiconformer phytochemicals library are shown in Table 2, along with representative calculated/predicted molecular, physicochemical and pharmacokinetic properties. These hits were docked into the catalytic site of PLpro using MOE 2022.02. A total of five poses of each of the hits were generated and further examined for their interactions with residues lining the PLpro catalytic site. Representative poses (hits **11** and **12,** Table 2), along with the 2D interaction diagrams are shown in Figures 5-6 (PDB ID 6WUU) and 7-8 (PDB ID 6WX4). The native ligands were docked in their respective crystal structures for optimizing the docking protocol settings. The docked poses and the 2D interaction diagrams of both the crystal structure ligands can be found in the *Supporting Information section* (Figures 1S and 2S). The corresponding docking scores are shown in Table 3.

**Figure 2.**
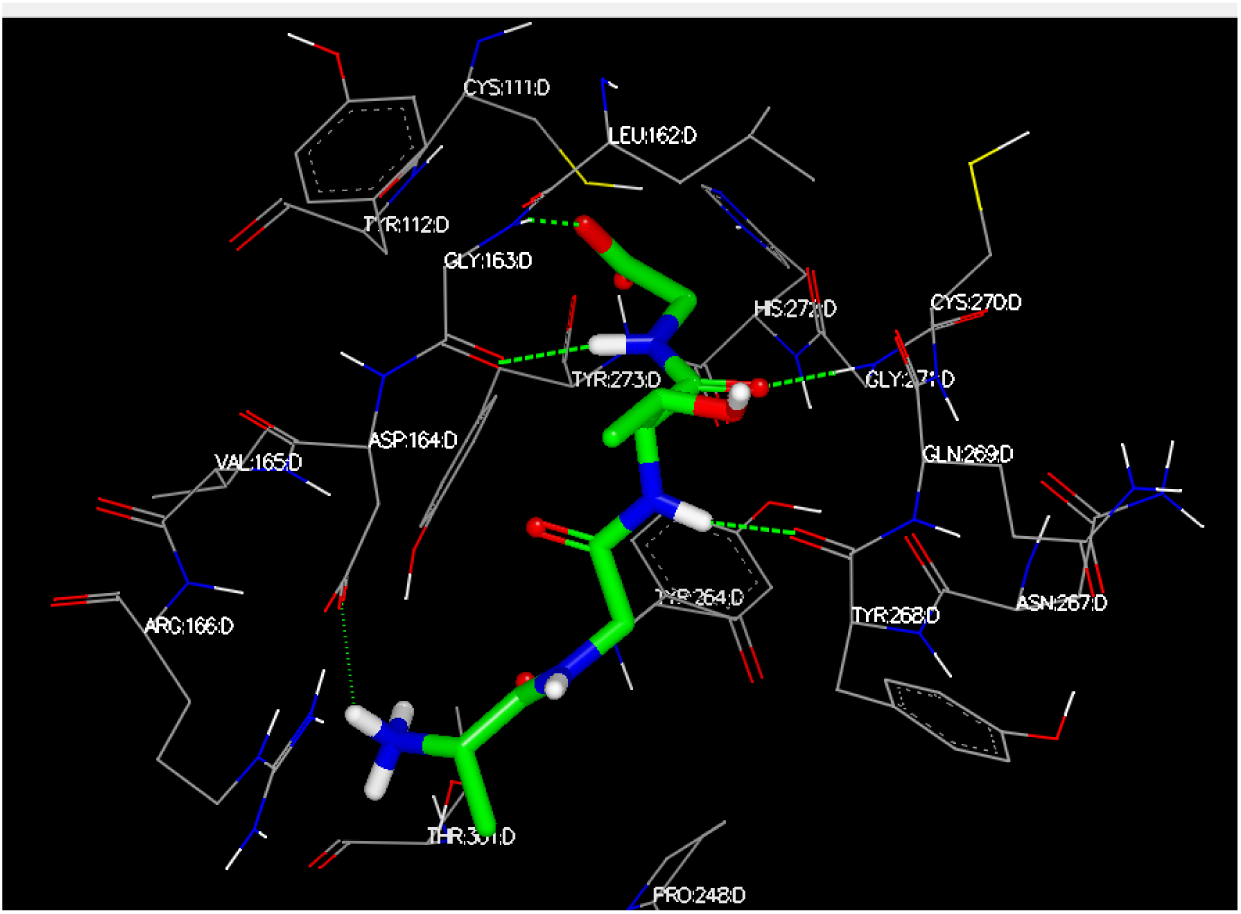
Representative top-scoring tetrapeptide 1 (Table 1) docked in the PLpro active site

**Figure 3.**
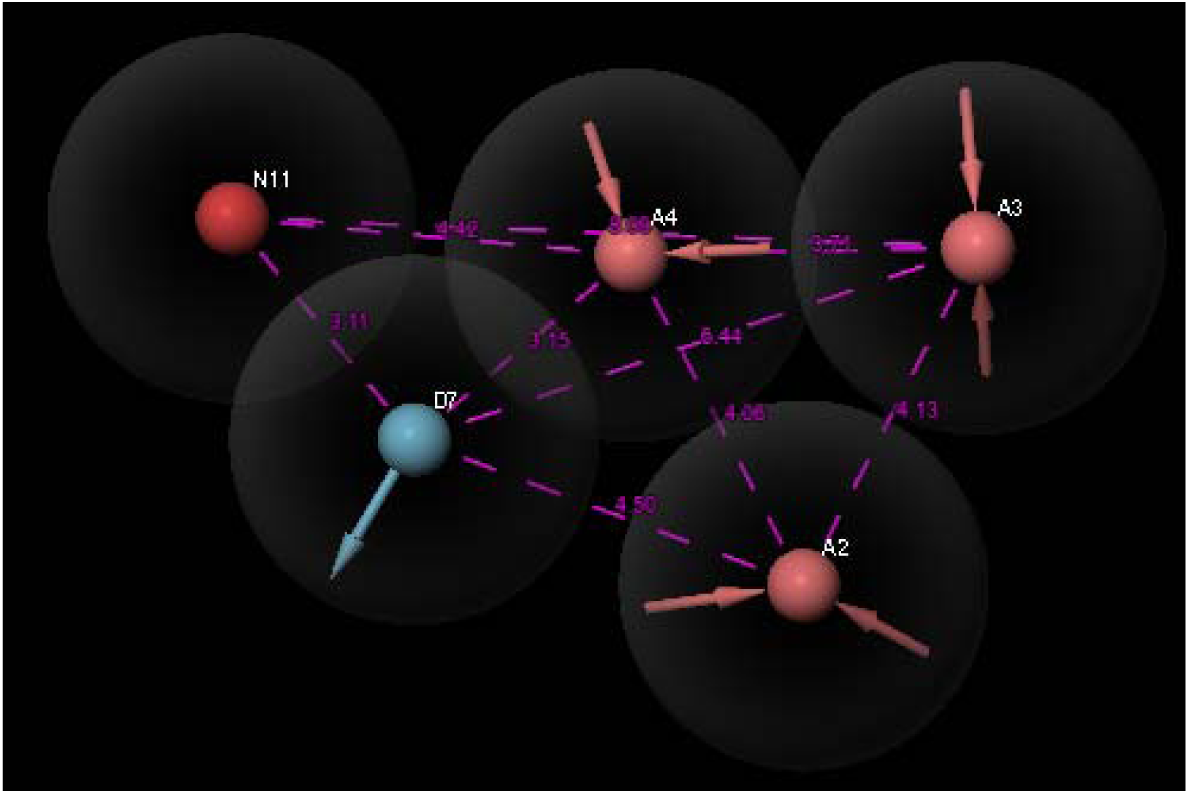
Top-scoring five-point pharmacophore (AAADN) with interfeature distances and relative arrangement

As seen from Figure 5, the two top-scoring poses of **11** in the active site of PLpro (PDB ID 6WUU) were flipped; in pose 1, the –CH_2_OH donor of **11** formed a H-bond with Asp164 and secondary *4’*–OH of the sugar moiety interacted with Tyr268 via H-bonding, while in pose 2, the *2’*-OH of **11** formed a H-bond with Asp164. Given the polar nature of the proximal and distal portions of **11**, the original and flipped poses did well in compensating the binding energy by forming multiple interactions with the active site residues. The salt-bridge interaction of the – COOH of **11** with nearby Arg166 was absent in both the top-scoring poses. This could be attributed to the fairly rigid framework of **11**. The situation was bit different for the other crystal structure of PLpro (PDB ID 6WX4). In Figure 7, both the top-scoring poses of **11** were found to be similar with subtle difference in the orientation of the terminal groups. Interestingly, the – COOH group of **11** formed a salt-bridge interaction with Lys157 (both poses). The polyol –OH groups interacted with PLpro active site residues with fewer H-bond interactions with Asp164, Thr301, and Tyr268. Overall, these multiple interactions led to slightly better docking score in the second crystal structure (PDB ID 6WX4). The docking scores of **11** were less than the corresponding scores of the native ligands of both the crystal structures (Table 3).

On similar lines as **11**, both the top-ranking poses of **12** showed flipped orientation in the active site of PLpro (PDB ID 6WUU) (Figure 6). The sugar moiety was observed to form couple of H-bonds with Met208 and Asp302 (pose 1), while the pose 2 formed a lone H-bond with Arg166. The propionyl moiety of pose 1 (*guache* conformation around C_2_-C_3_ bond) formed a salt-bridge interaction with Arg166 (Figure 6b); a similar interaction was absent in pose 2 (*anti* conformation around C_2_-C_3_ bond) (Figure 6c) due to flipping. In both poses, the propionyl moiety was exposed to solvent. Additionally, the benzofuran ring formed arene-H interaction with Asp164 only in pose 1 (Figure 6b). The docked poses of **12** in the PLpro structure (PDB ID 6WX4) (Figure 8) exhibited similar trend with respect to the flipped poses. While the propionyl side chain formed H-bonds with key residue, Cys111, the sugar moiety interacted with Thr301 via H-bond (Figure 8b). In pose 2, the sugar moiety formed extensive H-bond network with Cys111, Cys270, Gly 271 and Tyr268 and the propionyl group was involved in arene-H interaction with Tyr264 (Figure 8c). Overall, both poses 1 and 2 of **12** were distinct due to the moderate flexibility to the molecular structure. The docked poses of hits **13** to **15** in both crystal structures are given in *Supporting Information* section (Figures 3S to 8S).

Table 2 lists all the five hits, their structures and relevant information (CAS number, molecular formula and weight) along with the calculated/predicted molecular, physicochemical, pharmacokinetic and toxicity properties. The hits were all derived from originally phytochemical library. Due to the presence of significant proportion of heteroatoms in the molecular composition, the hits were appreciably polar, violated the Lipinski’s Rule of 5 filter and showed poor ADME predictions. This was no surprise given the nature of the substrate-binding site of the PLpro enzyme, evolved to identify peptides. The idea is to further transform the hits into more drug-like leads and optimize them along the way. The interesting part is that the hits were derived from a novel strategy, i.e., peptide mold, which can potentially be applied to any protein target, leave aside the proteases such as PLpro. The first hit, **11** (Geniposidic acid, Table 2) belongs to iridoid glycoside class, found in significant number of plants. It possesses multiple pharmacological activities, with main activity being anticancer.^61^ A recent article reports its antiviral activity against white spot syndrome virus via reducing *STAT* (signal transducer and activator of transcription) gene expression, thereby reducing viral replication.^62^ Compound **11** has been studied as tool molecule for white spot syndrome virus replication inhibition. The second hit, **12**, is a ZINC database member (ZINC39010597)^63a^, which has not been tested in any assay. The activity predictions using SwissTargetPrediction^64^, indicated that it was likely to be a transporter inhibitor, without any peculiar details on the transporter.

Compound **13** (Table 2), a dihydroxybenzamide derivative of Ala-Ser dipeptide, is present in ZINC database (ZINC2356677764)^63b^ and has not been evaluated in any in vitro biological assay. The target prediction for **13** using SwissTargetPrediction^64^ yielded caspase-1 as the top-raking target. Further literature searches in this direction led to an interesting fact that few viruses are dependent on the host caspases for proteolytic processing of the viral components.^65^ A caspase inhibitor is likely to demonstrate antiviral effects. On the other hand, caspases are Zn^2+^-dependent proteolytic enzymes.^66^ The N-terminal modified Ala-Ser, i.e., **13**, is likely to bind in the active site of caspases. Inhibition of host caspases by a potential ligand may disrupt virus particle processing. In short, **13**, a virtual screening hit, can be a potential PLpro inhibitor by virtue of its pharmacophoric features, and it may also hit other proteolytic enzymes, such as caspases, which in turn, may indirectly alter viral spread and infection. The cyclic pentapeptide, **14** (WS-7338A), is unique due to its structural features. It is reported to exhibit potent endothelin receptor antagonist activity.^67^ A recent report discussed the potential utility of endothelin receptor antagonist as antiviral entities.^68^ The hit **15** has been reported as dipeptidyl aminopeptidase IV (DPP IV, EC 3.4.14.5) inhibitor long ago.^69^ Recently, it has been argued whether DPP IV inhibitors could be used for the development of novel viral entry inhibitors for emerging viral infections.^70^ To summarize, the hits from virtual screening of phytochemicals library have a potential to exhibit antiviral action, most likely, via PLpro inhibition. The possibility of these agents to hit other proteolytic enzymes is likely to exist due to their peptidic (**13**-**15**) or non-peptidic (**11** and **12**) nature. All the five high-quality hits have originally been screened during pharmacophore search, followed by molecular docking.

**Figure 4.**
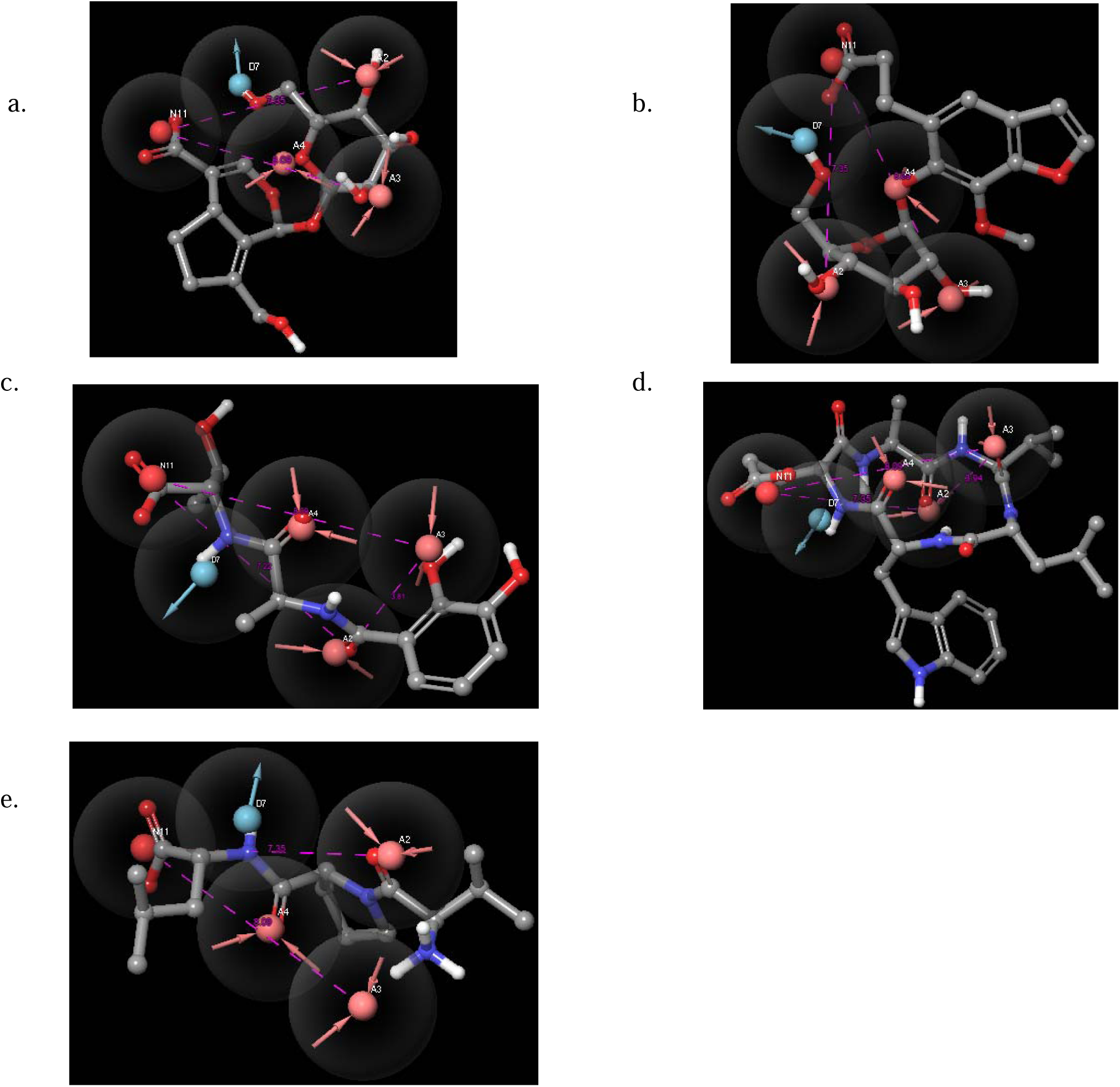
Mapping of pharmacophoric features back onto the hits from multiconformer phytochemicals library - a) Geniposidic acid (**11**), b) ZINC39010597 (**12**), c) ZINC2356677764 (**13**), d) WS-7338A (**14**) and e) Diprotin A (**15**). The directional nature of HDon and HAcc features are clearly seen in the provided figures.

**Figure 5.**
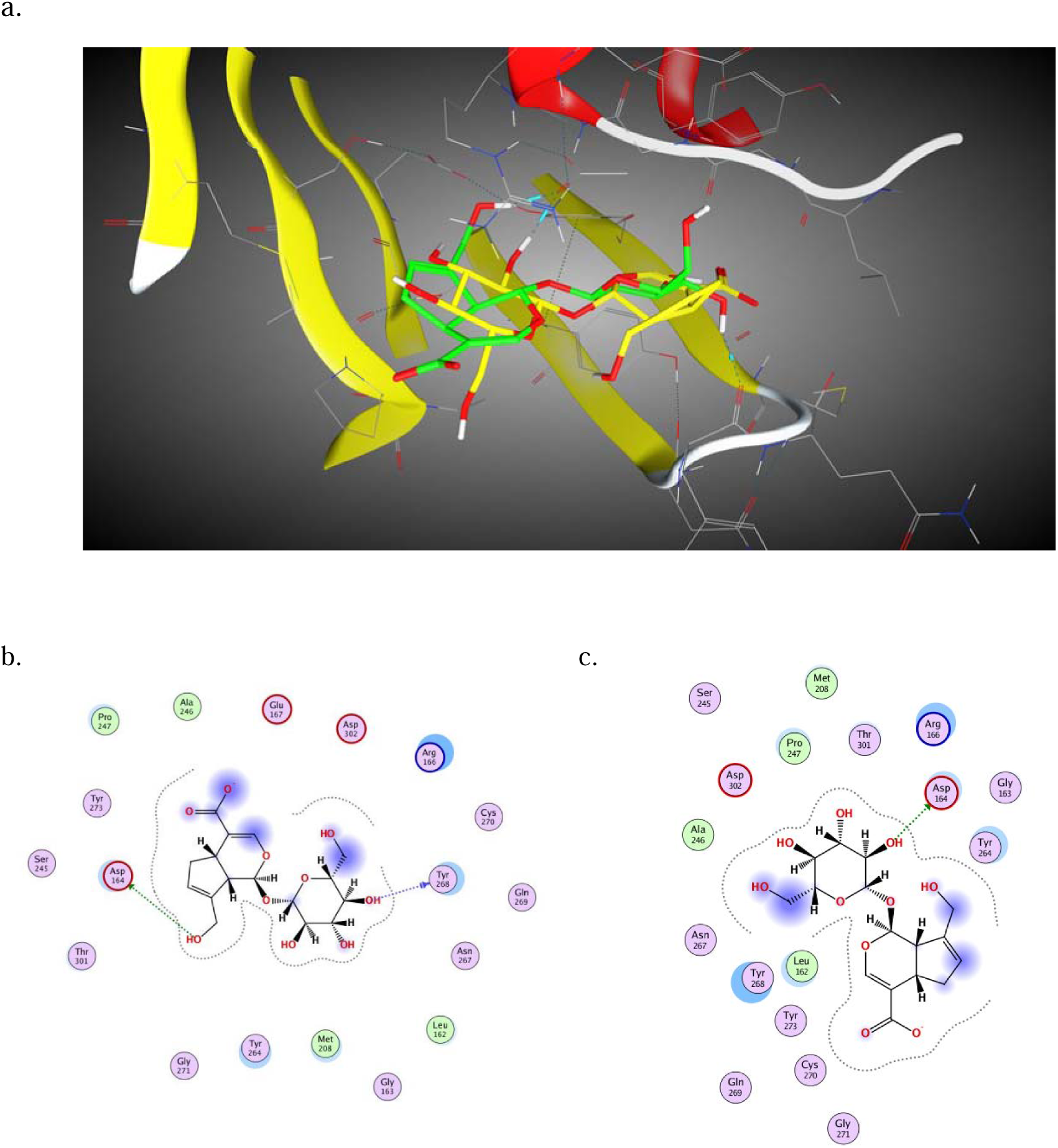
Docked poses 1 (green capped-stick representation) and 2 (yellow capped stick Representation) of 11 in the SARS-CoV-2 PLpro catalytic site (PDB ID 6WUU); 2D interaction diagram of b. 11 – Pose 1 and c. 11 – Pose 2

**Figure 6.**
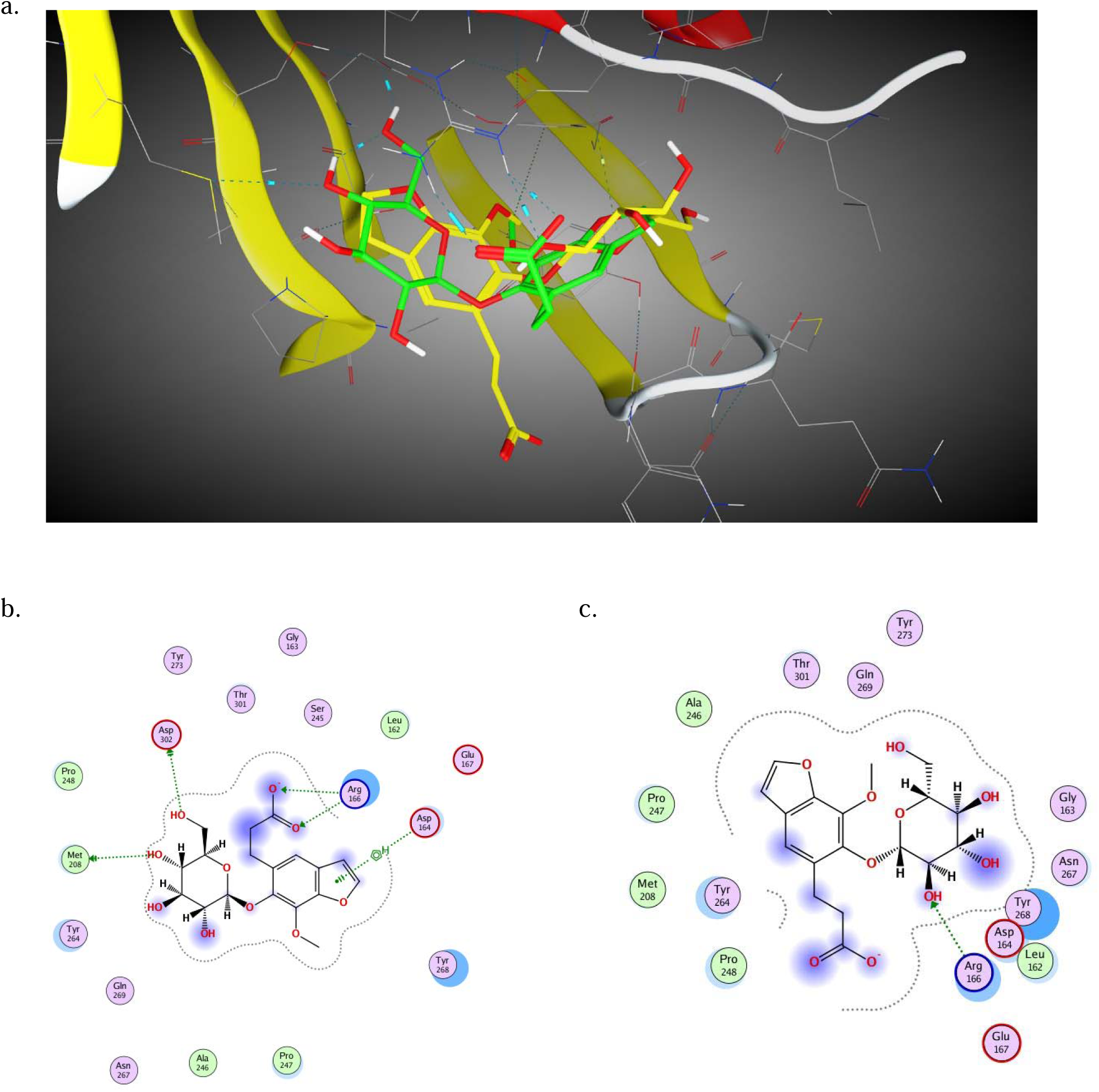
a. Docked poses 1 (green capped-stick representation) and 2 (yellow capped stick Representation) of 12 in the SARS-CoV-2 PLpro catalytic site (PDB ID 6WUU); 2D interaction diagram of b. 12 – Pose 1 and c. 12 – Pose 2

**Figure 7.**
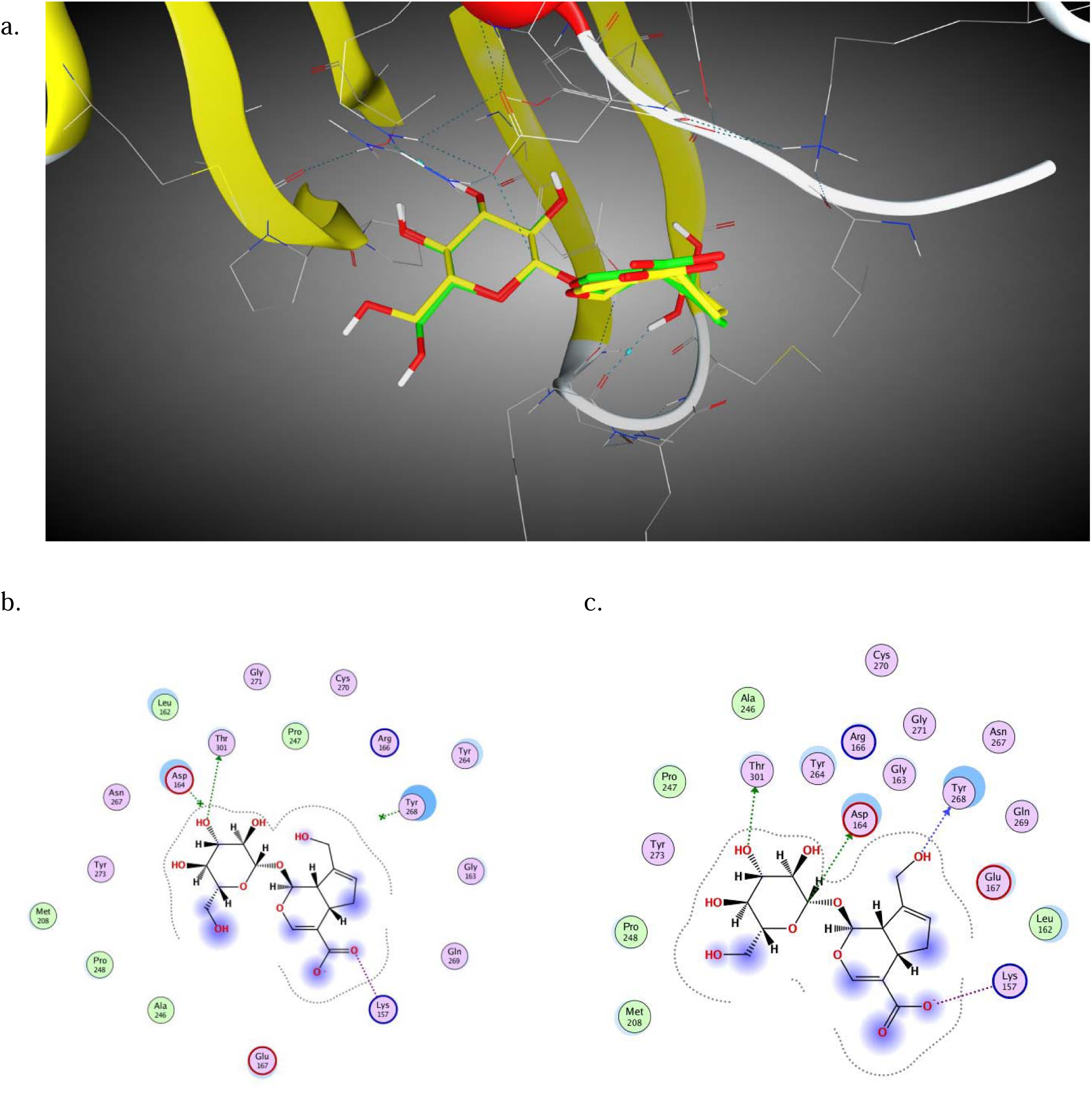
a. Docked poses 1 (green capped-stick representation) and 2 (yellow capped stick Representation) of 11 in the SARS-CoV-2 PLpro catalytic site (PDB ID 6WX4); 2D interaction diagram of b. 11 – Pose 1 and c. 11 – Pose 2

**Figure 8.**
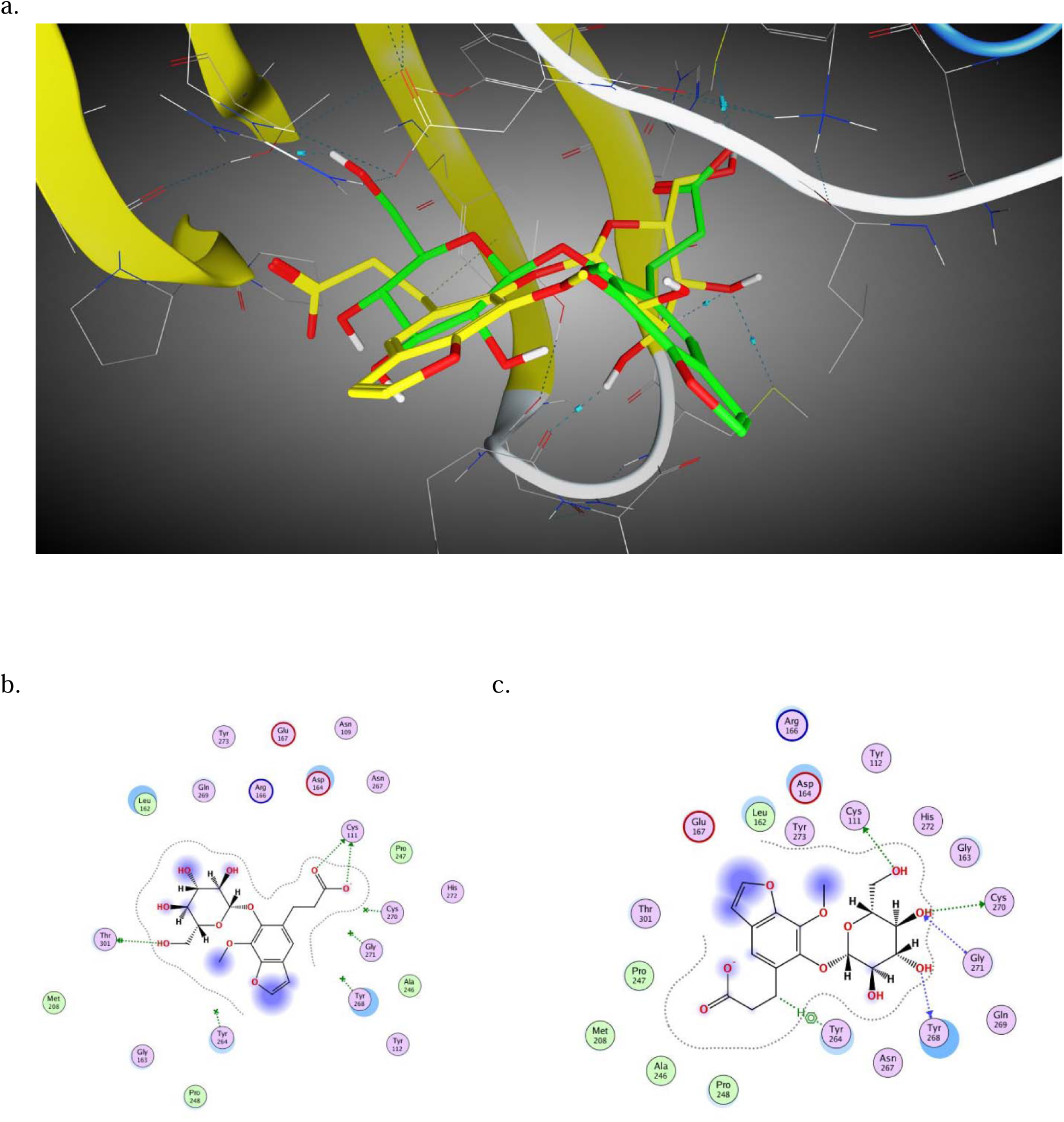
a. Docked poses 1 (green capped-stick representation) and 2 (yellow capped stick Representation) of 12 in the SARS-CoV-2 PLpro catalytic site (PDB ID 6WX4); 2D interaction diagram of b. 12 – Pose 1 and c. 12 – Pose 2

**Table 2.**
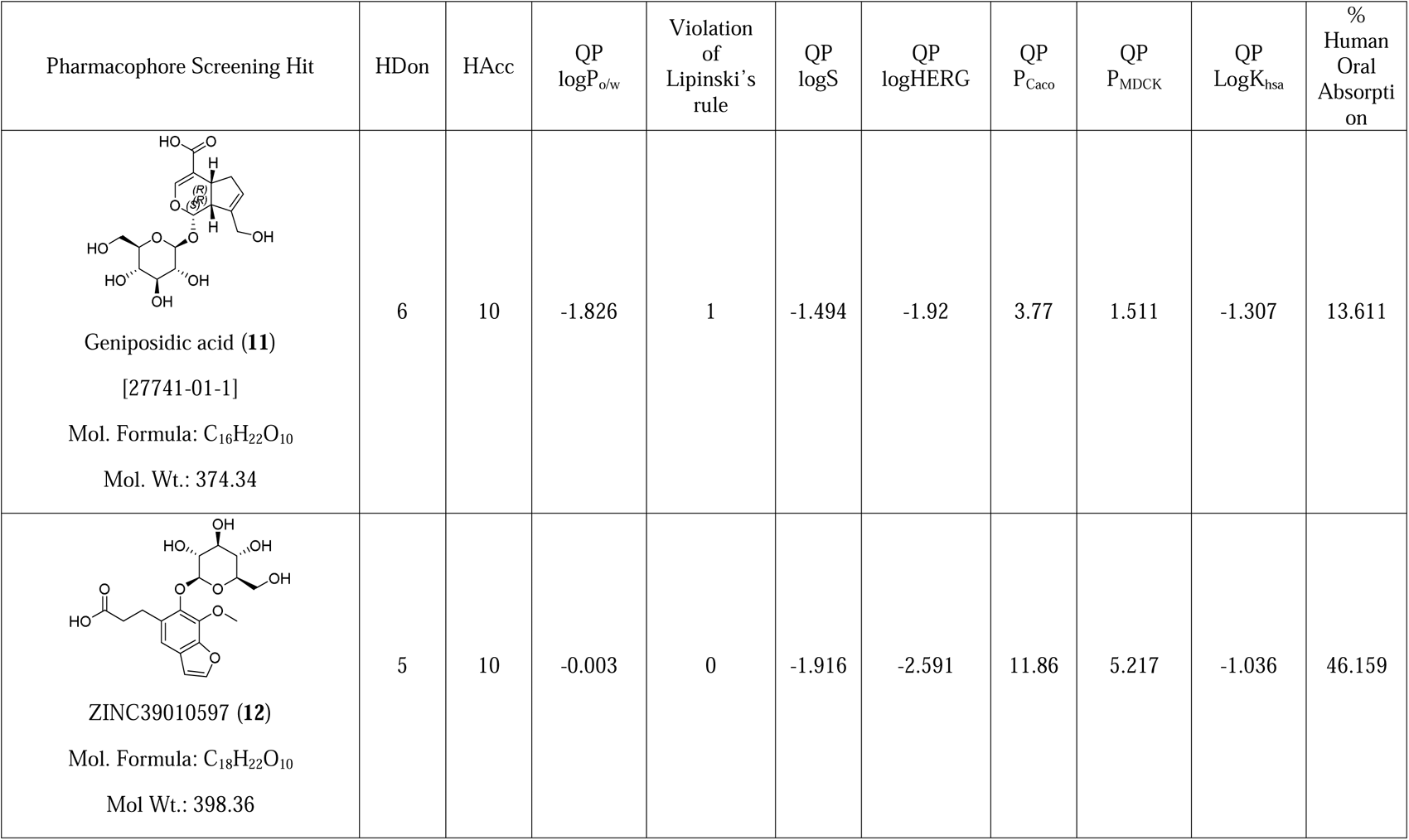
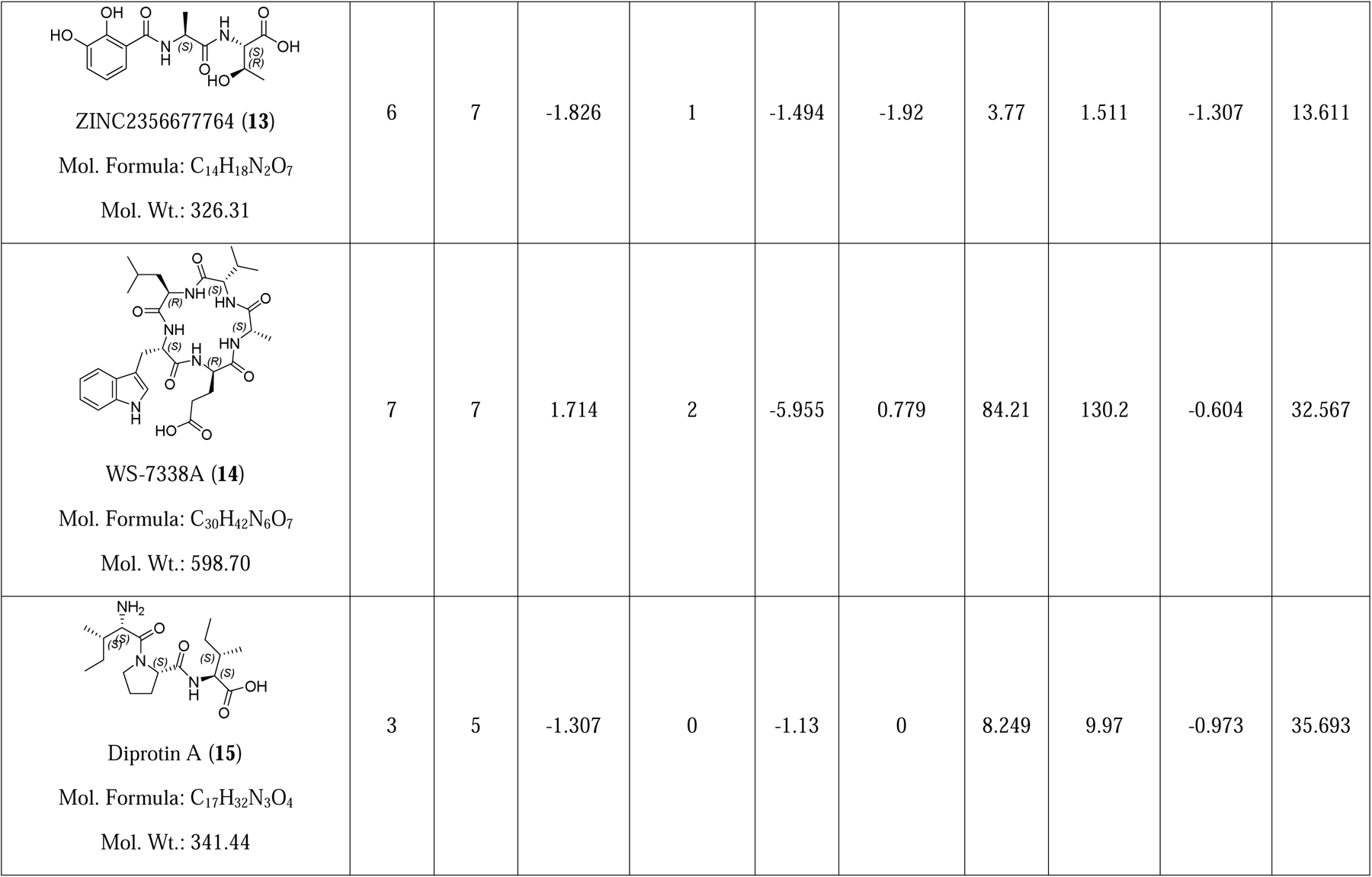
Calculated/predicted molecular, physicochemical and pharmacokinetic properties of top-five hits from pharmacophore screening.

| Pharmacophore Screening Hit | HDon | HAcc | QP<br>logP<sub>o/w</sub> | Violation<br>of<br>Lipinski's<br>rule | QP<br>logS | QP<br>logHERG | QP<br>P<sub>Caco</sub> | QP<br>P<sub>MDCK</sub> | QP<br>LogK<sub>hsa</sub> | %<br>Human<br>Oral<br>Absorption |
| --- | --- | --- | --- | --- | --- | --- | --- | --- | --- | --- |
| **Geniposidic acid (11)**<br>[27741-01-1]<br>Mol. Formula: C<sub>16</sub>H<sub>22</sub>O<sub>10</sub><br>Mol. Wt.: 374.34 | 6 | 10 | -1.826 | 1 | -1.494 | -1.92 | 3.77 | 1.511 | -1.307 | 13.611 |
| **ZINC39010597 (12)**<br>Mol. Formula: C<sub>18</sub>H<sub>22</sub>O<sub>10</sub><br>Mol Wt.: 398.36 | 5 | 10 | -0.003 | 0 | -1.916 | -2.591 | 11.86 | 5.217 | -1.036 | 46.159 |
| **ZINC2356677764 (13)**<br>Mol. Formula: C<sub>14</sub>H<sub>18</sub>N<sub>2</sub>O<sub>7</sub><br>Mol. Wt.: 326.31 | 6 | 7 | -1.826 | 1 | -1.494 | -1.92 | 3.77 | 1.511 | -1.307 | 13.611 |
| **WS-7338A (14)**<br>Mol. Formula: C<sub>30</sub>H<sub>42</sub>N<sub>6</sub>O<sub>7</sub><br>Mol. Wt.: 598.70 | 7 | 7 | 1.714 | 2 | -5.955 | 0.779 | 84.21 | 130.2 | -0.604 | 32.567 |
| **Diprotin A (15)**<br>Mol. Formula: C<sub>17</sub>H<sub>32</sub>N<sub>3</sub>O<sub>4</sub><br>Mol. Wt.: 341.44 | 3 | 5 | -1.307 | 0 | -1.13 | 0 | 8.249 | 9.97 | -0.973 | 35.693 |

**Table 3.** Docking score of the five hits from pharmacophore screening along with the crystal structure ligands.

| Pharmacophore Screening Hit/Crystal structure ligand<sup>a</sup> | Docking score (PDB ID 6WUU) | Docking score (PDB ID 6WX4) |
| --- | --- | --- |
| 6WUU ligand | -10.64 | ND |
| 6WX4 ligand | ND | -9.78 |
| **11** | -7.15 | -7.16 |
| **12** | -7.56 | -7.47 |
| **13** | -6.82 | -7.51 |
| **14** | -7.53 | -8.14 |
| **15** | -7.62 | -7.43 |

<sup>a</sup>Docking score of the top-ranking pose

To obtain better insights into the interactions between **13** and **15** with SARS-CoV-2 PLpro, we performed all-atom molecular dynamics simulations in an explicit water box. Furthermore, the stability of the protein ligand was estimated using an end-state free-energy method like MM-GB/SA on the MD sampled conformations. We used the backbone RMSD to understand the progression of the MD simulations and backbone RMSF for the overall dynamics of the protein-ligand complexes. The SARS-CoV-2 PLpro protein was found to be very flexible as shown by the backbone RMSD (**Figure 9A**) and the backbone RMSF (**Figure 9C**). Upon closer inspection, we observed that large fluctuations were observed in the C-terminal region from amino acid numbers 170-320. During the simulation, the protein significantly deviates from the starting conformation until 100 ns. The normalized frequency distribution shows that a large population of the protein conformations fluctuates around RMSD between 2-3 Å which indicates significant deviation from the starting geometry. We hypothesize that it could be attributed to the ligand binding wherein the protein samples conformational states are distinct from the starting geometry. To understand the strength of the binding of the ligands, we used the MMGBSA method to compute the binding energies, due to computational cost we excluded the entropy calculations, and therefore, the binding energies we obtained shall be treated as effective binding energies (ΔG_eff_). Our MD simulations suggest that **15** binds with far superior binding affinity than **13** (**Figure 9B**). Furthermore, we observed several unbinding events for **13**, as a result, the mean ΔG_eff_ for **13 is -22.61 kcal/mol** less than **15**. Compound **15** is well suited to occupy the SARS-CoV-2 PLpro binding site as the proline at the centre of the **15** creates a bend in the backbone that enhances the binding stability. This structural feature is absent in **13** which may have caused poor affinity towards SARS-CoV-2 PLpro.

Figure 10 shows the docked pose of **15** following MD simulations along with the 2D interaction diagram and the interaction surface for clear visibility of the substrate-binding site of PLpro. As seen in Figure 5S (Supporting Information section) and Figure 10, the interactions of **15** with binding site residues are conserved. The amino terminal group formed electrostatic interactions with Asp164 and Asp302, along with H-bonding interactions with Thr301. The other electrostatic interaction of anionic carboxylate of **15** with Arg166 (Figure 5S) was absent in the post-MD simulation pose (Figure 10). A new H-bond was formed between C-terminal –C=O of **15** and Tyr273 (Figure 10) which was not observed in the pre-MD pose (Figure 5S).

Additionally, one of the methylene groups of pyrrolidine ring of 15 interacted with Tyr264 via arene-H interaction (Figure 10). The superposed active site residues of the substrate-binding site residues along with the pre- and post-MD simulations are shown in Figure 11 for comparison. As seen from Figure 11, the MD simulations led to moderate changes in the backbone and side-chain atoms of the substrate-binding site residues, thereby optimizing the interactions with 15, which were stable over the simulation intervals, giving rise of gain in binding energy.

**Figure 9.**
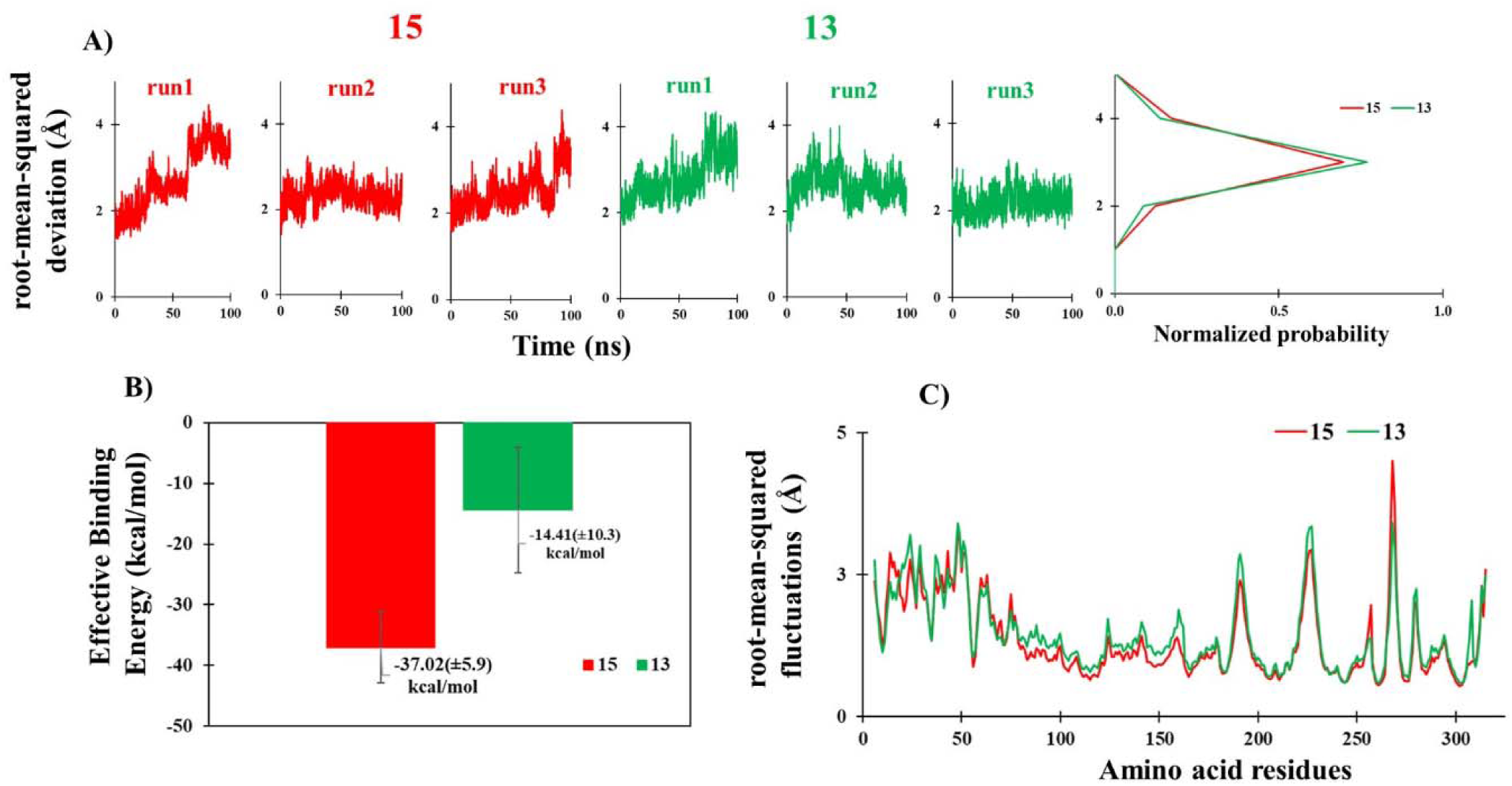
A. Time evolution plot of backbone atom RMSD for **15** complexed with SARS-CoV-2 PLpro (red traces) and **13** complexed with SARS-CoV-2 PLpro (green traces); B. Bar diagram depicting the effective binding energy of **15** (red bar) and **13** (green bar) with SARS-CoV-2 PLpro; C. The root-mean-squared fluctuations of the backbone atoms SARS-CoV-2 PLpro when bound to **15** (red trace) and **13** (green trace).

**Figure 10.**
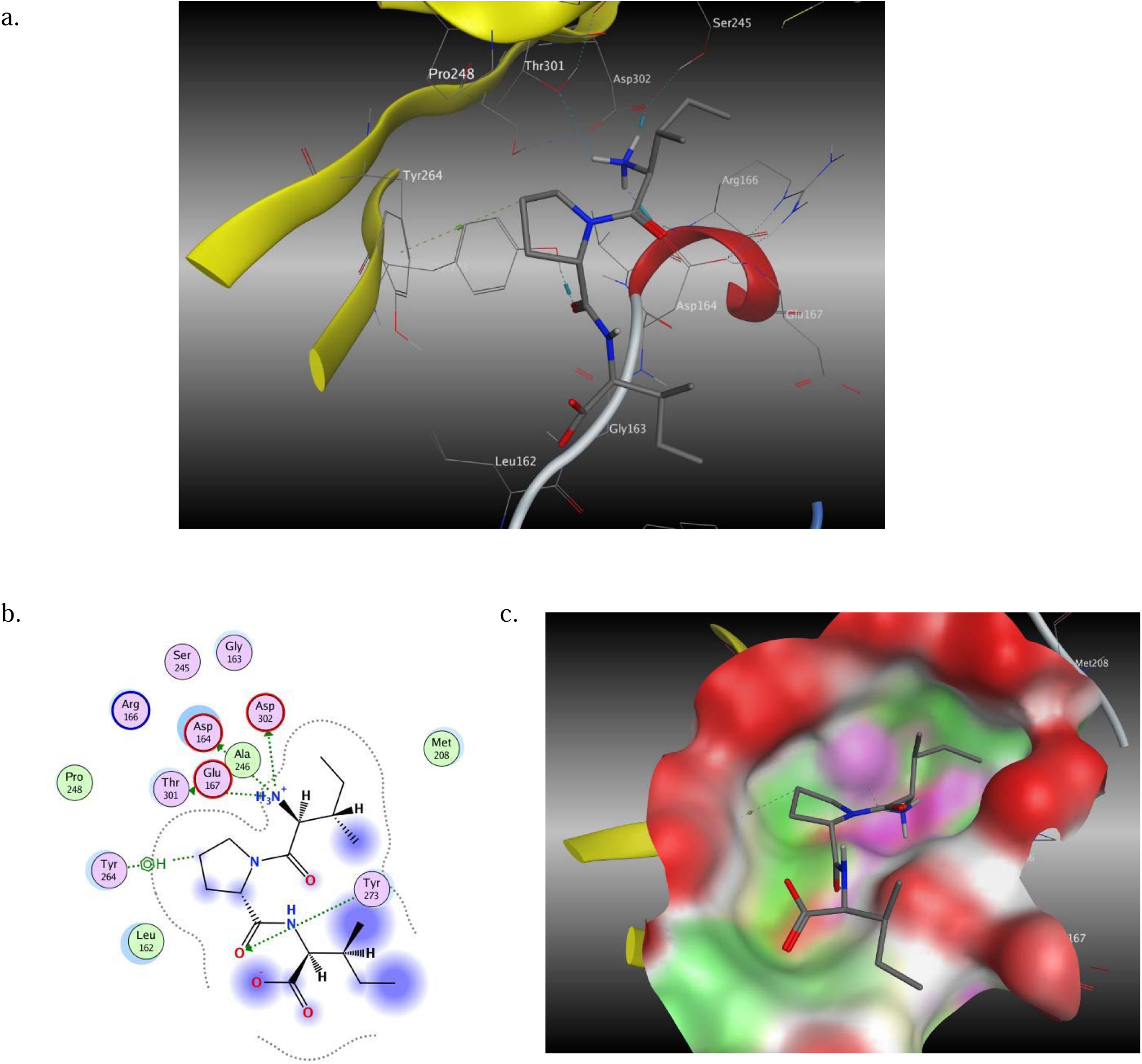
a. Docked pose of hit **15** (capped-stick representation) following MD simulations; b. 2D interaction diagram of 15 in the substrate-binding site of PLpro; c. Interaction surface of 15 with binding site residues (green: hydrophobic, pink: polar, red: exposed)

**Figure 11.**
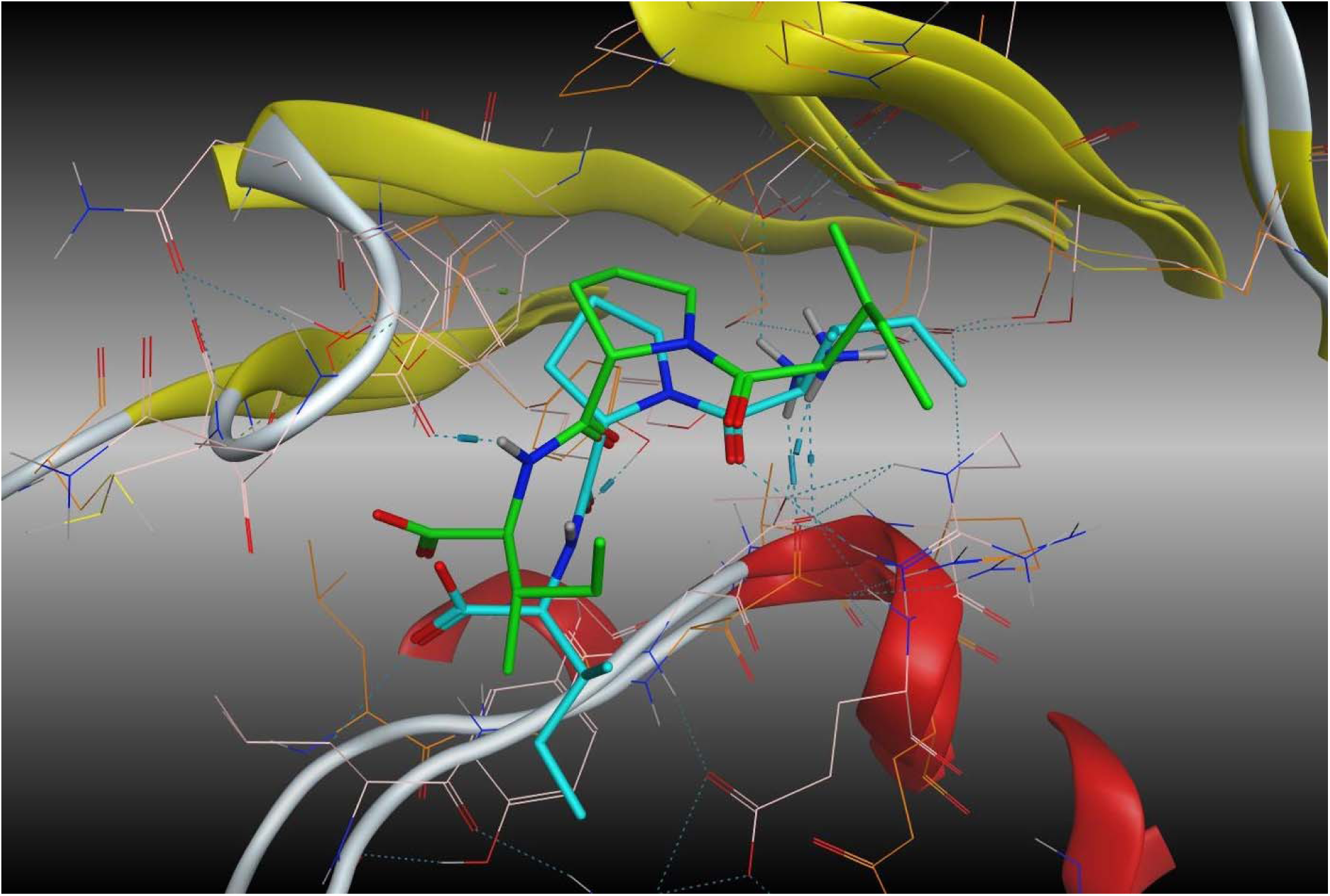
Superposed view of SARS-CoV-2 PLpro substrate-binding site pre-(light-pink line representation) and post-MD (orange line representation) simulations. Hit 15 pre-MD (green capped-stick representation) and post-MD (cyan capped-stick representation) simulations are shown for comparison

The direct docking of the phytochemicals library in the substrate-binding site SARS-CoV-2 PLpro (PDB ID 6WX4) and the subsequent binding free energy estimations were performed to compare the utility of the peptide mold strategy over the conventional structure-based methodology. The corresponding results for the top-five hits are shown in Table 4. Hits **16** and **20** were amphoteric, hits **17** and **19** were anionic and hit 18 was neutral. Their molecular weights were >500 Da, except **16**. Since the hits were arranged in the ascending order of binding free energy values, it was no surprise that the bigger molecules yielded higher binding free energy estimates. These hits were docked in similar manner as hits from pharmacophore screening in the substrate binding site of PLpro (PDB IDs 6WUU and 6WX4) and compared. The results are listed in Table 5. The top-ranking docked poses 1 and 2 along with their 2D interaction diagrams of hit **18** in the substrate-binding site of PLpro (PDB ID 6WX4) are shown for comparison in Figure 10. Table 5 also lists the H-bonds formed between the top-ranking pose of the hits and PLpro substrate-binding site residue. The corresponding docked poses and the 2D interaction diagrams of hits **16**, **17**, **19** and **20** in PLpro site (PDB ID 6WX4) are given in *Supporting Information* section (Figures 9S to 12S).

Hit **18** formed three H-bonds (pose 1) with Gly163, Gly271 and Cys270 (Figure 10), while in pose 2, additional H-bond with Tyr268 was observed. Since **18** lacked anionic group (unlike other hits), no salt bridge interaction with PLpro cationic residues was seen. The tetrahydrofuro[2,3-*d*][1,3]dioxol-6-ol moiety was exposed to solvent at the proximal end of the substrate-binding site of the enzyme. Both the poses exhibited similar binding modes with subtle differences in the orientation of the benzyl group. No information of **18** could be retrieved from literature except the CAS No.: 1191113-39-9. It has not been previously used for any purpose at all. Hit **16** (CAS No. 455325-25-4) was reported in one research paper two decades ago.^71^ Hit **17** and **20** were not reported in the chemical literature at all. Though the CAS No. was available for hit **19** (1241902-07-7), no biological evaluation was ever carried out on it.

**Table 4.**
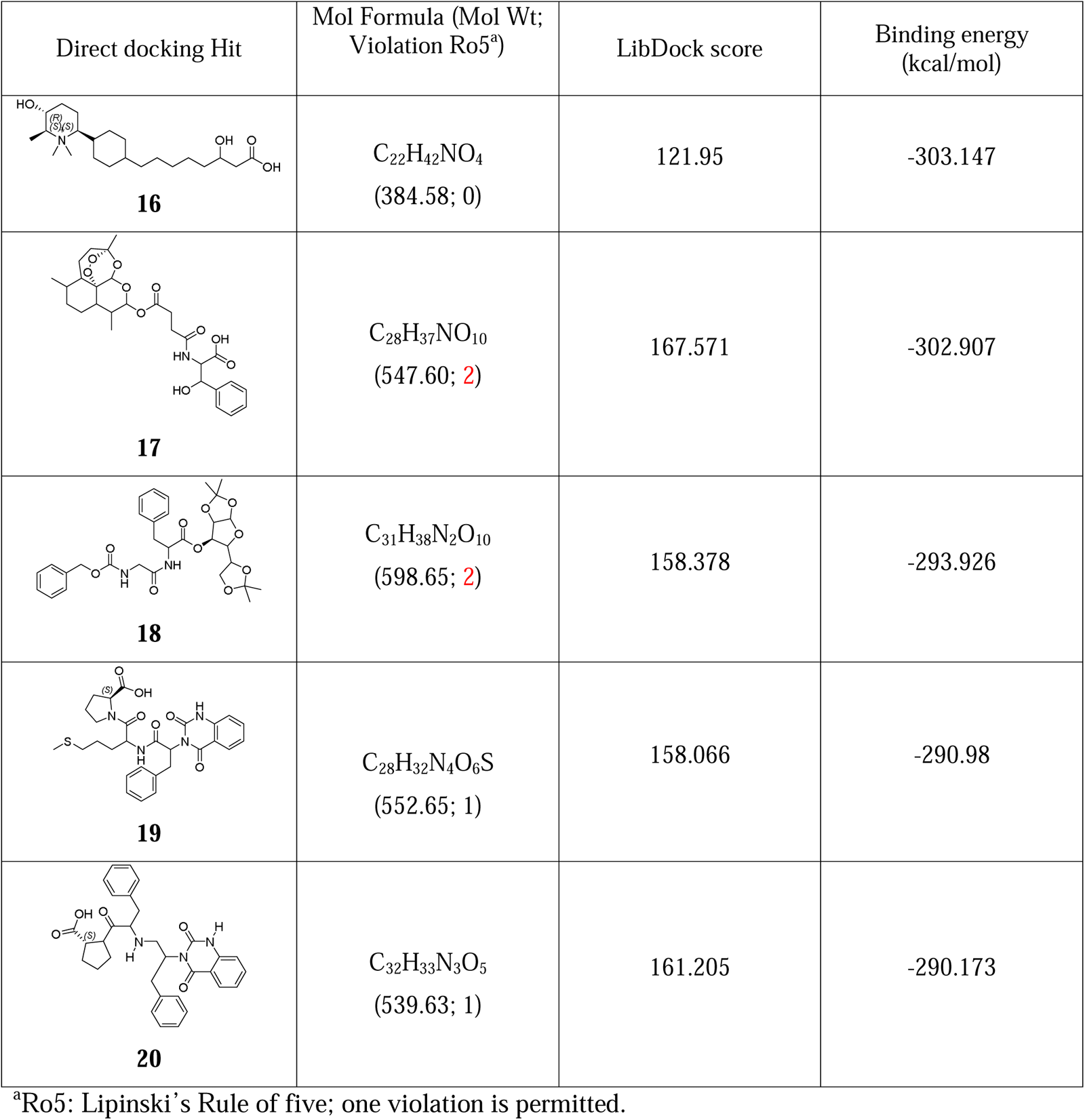
Docking scores and the binding free energy estimates of the five hits from direct docking of the phytochemicals library in the susbtrate-binding site of PLpro (PDB ID 6WX4)

| Direct docking Hit | Mol Formula (Mol Wt;<br>Violation Ro5<sup>a</sup>) | LibDock score | Binding energy<br>(kcal/mol) |
| --- | --- | --- | --- |
| **16** | $C_{22}H_{42}NO_4$<br>(384.58; 0) | 121.95 | -303.147 |
| **17** | $C_{28}H_{37}NO_{10}$<br>(547.60; 2) | 167.571 | -302.907 |
| **18** | $C_{31}H_{38}N_2O_{10}$<br>(598.65; 2) | 158.378 | -293.926 |
| **19** | $C_{28}H_{32}N_4O_6S$<br>(552.65; 1) | 158.066 | -290.98 |
| **20** | $C_{32}H_{33}N_3O_5$<br>(539.63; 1) | 161.205 | -290.173 |

<sup>a</sup>Ro5: Lipinski's Rule of five; one violation is permitted.

**Table 5.**
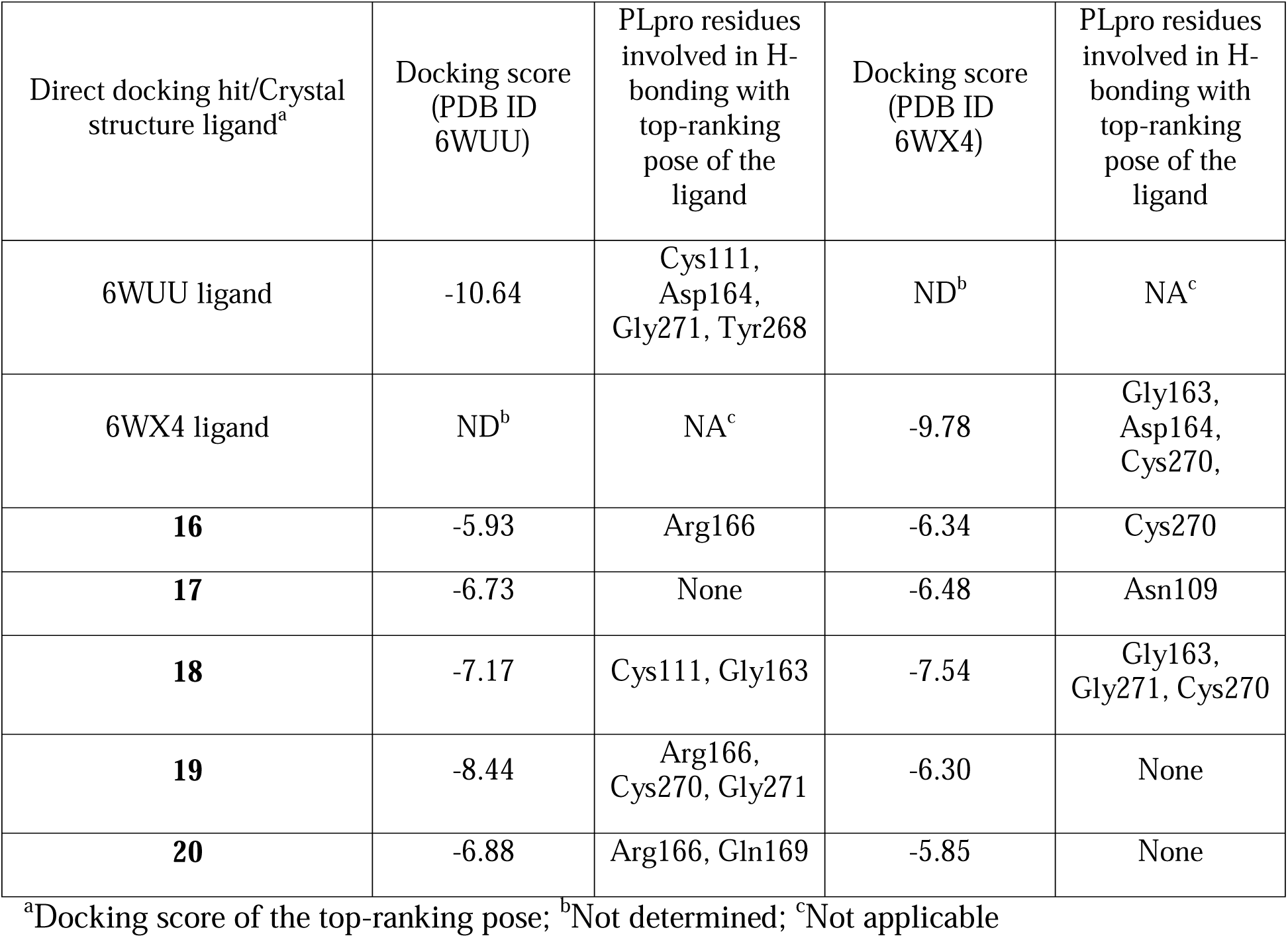
Docking score of the five hits from direct docking of phytochemicals library along with the crystal structure ligands.

Overall, the top-scoring hits scouted using the direct docking from the same phytochemical library were observed to be bigger and less drug-like, compared to the hits obtained using the complete process flow based on peptide mold concept. When the overlap of top-500 hits from both the peptide mold and direct docking strategies was investigated, we found less than 50 hits common both workflows (data not shown).

**Figure 11.**
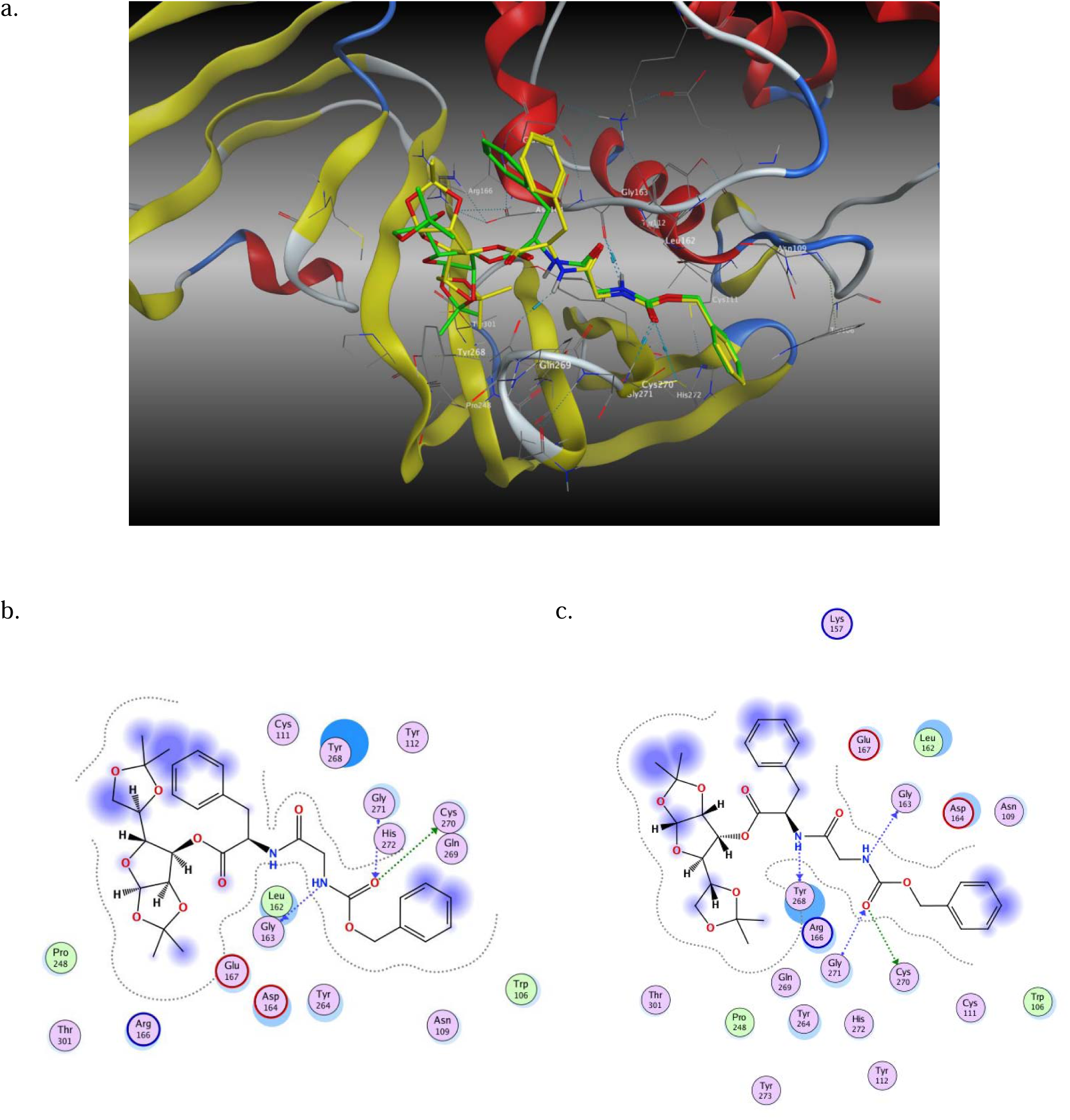
a. Docked poses 1 (green capped-stick representation) and 2 (yellow capped stick Representation) of 18 in the SARS-CoV-2 PLpro catalytic site (PDB ID 6WX4); 2D interaction diagram of b. 18 – Pose 1 and c. 18 – Pose 2

## Conclusions

A novel concept, ‘peptide mold’ based on the use of tetrapeptides for creating an imprint of steric and electrostatic features of the substrate-binding site of SARS-CoV-2 PLpro is presented. The created imprint comprising of various drug-receptor interaction features and the shape of the catalytic site was further used for developing a five-point pharmacophore model, which in turn, was used for ligand-based virtual screening of a phytochemicals library. The resulting high-quality hits were further docked multiple times to fine-tune the selection of promising small-molecule phytochemical hits. The extended molecular dynamics simulations of the top-two hits from final docking runs were used for further calculation of binding energy values and for checking the stability of binding. SARS-CoV-2 PLpro was used a test case for exploring the potential of this approach. The conventional direct docking approach was used for virtual screening of the hits to compare both the strategies. The resultant hits were bigger in size and less drug-like with little overlap between the hits from both workflows. The presented concept has not been reported in the literature so far. It can be extended to any protein for catalytic, substrate-binding and/or allosteric binding site and has enormous potential to make virtual screening more productive with quality hits. This, in turn, can increase the output and success of the experimental medium- and high-throughput screens. Further exploration of the peptide mold strategy by the scientific community will definitely lead to more and more success stories.

## ASSOCIATED CONTENT

### Supporting Information

The docked poses of the crystal structure ligands (PDB IDs 6WUU and 6WX4) and the hits from peptide mold strategy and direct docking in the substrate-binding pocket of SARS-CoV-2 PLpro and their corresponding 2D interaction diagrams are provided. This material is free of charge.

## AUTHOR INFORMATION

### Author Contributions

PSK conceived and articulated the concept and other authors contributed to further development of the presented work. The manuscript was written through contributions of all authors. All authors have given approval to the final version of the manuscript.

### Notes

The presented work was culmination of concerted efforts of the team during COVID-19 pandemic. The work was part of Drug Discovery Hackathon 2020 (DDH2020) Challenge: Track 1: Drug design for anti-COVID-19 hit/lead molecule generation. The work was shortlisted in Round 1. The authors declare no competing financial interest.

## ACKNOWLEDGMENT

The authors are thankful to OpenEye, Cadence Molecular Sciences, Schrödinger Inc., Dassault Systèmes, Organizers of Drug Discovery Hackathon 2020, for providing the software licenses to Prof. Prashant S. Kharkar’s laboratory, Institute of Chemical Technology (ICT), Mumbai. PSK thanks Prof. A. B. Pandit, Vice Chancellor, Institute of Chemical Technology, Mumbai, for providing necessary facilities.

## ABBREVIATIONS

SARS-CoV-2: Severe Acute Respiratory Syndrome Coronavirus 2
PLpro: Papain-like Protease
WHO: World Health Organization
COVID-19: Coronavirus Disease 2019
mRNA: Messenger RNA
Nsp: Non-structural Protein
HIV: Human Immunodeficiency Virus
PX12: Protein Disulfide Isomerase Inhibitor
NCEs: New Chemical Entities
CDAC: Centre for Development of Advanced Computing
AM1Bcc: Austin Model 1/Burkatzki-Cedillo-Cummins
MOE: Molecular Operating Environment
MACCS: Molecular Access System
SMILES: Simplified Molecular Input Line Entry System
OPLS3: Optimized Potentials for Liquid Simulations 3
GBVI: Generalized Born Volume Integral
WSA dG: Water Solvation Analysis Free Energy Difference
AMBER: Assisted Model Building with Energy Refinement
ZAFF: Zinc Amber Force Field
GAFF2: General Amber Force Field 2.0
TIP3P: Transferable Intermolecular Potential 3-Point
XP: Extra Precision
ADME/T: Absorption, Distribution, Metabolism, Excretion, and Toxicity
MD: Molecular Dynamics
NVT: Number of Particles, Constant Volume, and Constant Temperature
GPU: Graphics Processing Unit
PMEMD: Particle Mesh Ewald Molecular Dynamics
SHAKE: Scaled Harmonic Algorithm for Kinetic Energy
CPPTRAJ: C++ Trajectory Analysis Tool
MMGBSA: Molecular Mechanics Generalized Born Surface Area
EMM: Empirical Molecular Mechanics
GNP: Genetic Neighbourhood Potential
GGB: Generalized Gradient-Bond
GB: Generalized Born
GBOBC: Generalized Born Outer Bound
RMSD: Root Mean Square Deviation
Å: Angstroms
STAT: Signal Transducer and Activator of Transcription
QPLogPo/w: Predicted LogP Octanol/Water Partition Coefficient
QPLogS: Predicted LogS Solubility
QP LogHERG: Predicted Log HERG Channel Inhibition
QPPCaco: Predicted Permeability in Caco-2 Cell Monolayers
QPMDCK: Predicted Permeability in MDCK Cell Monolayers
QPKhsa: Predicted Human Serum Albumin Binding Affinity

