## Supporting Information for "Peptide Mold: A Novel Strategy for Mapping Potential Binding Sites in Protein Targets"

**Table of Contents**

| **Sr. No.** | **Description** | **Page No.** |
| --- | --- | --- |
| 1 | **Figure 1S.** Docked poses of crystal structure ligand (PDB ID 6WUU) | 3 |
| 2 | **Figure 2S.** Docked poses of crystal structure ligand (PDB ID 6WX4) | 4 |
| 3 | **Figure 3S.** Docked poses 1 and 2 of **13** in the SARS-CoV-2 PLpro catalytic site (PDB ID 6WUU) | 5 |
| 4 | **Figure 4S.** Docked poses 1 and 2 of **14** in the SARS-CoV-2 PLpro catalytic site (PDB ID 6WUU) | 6 |
| 5 | **Figure 5S.** Docked poses 1 and 2 of **15** in the SARS-CoV-2 PLpro catalytic site (PDB ID 6WUU) | 7 |
| 6 | **Figure 6S.** Docked poses 1 and 2 of **13** in the SARS-CoV-2 PLpro catalytic site (PDB ID 6WX4) | 8 |
| 7 | **Figure 7S.** Docked poses 1 and 2 of **14** in the SARS-CoV-2 PLpro catalytic site (PDB ID 6WX4) | 9 |
| 8 | **Figure 8S.** Docked poses 1 and 2 of **15** in the SARS-CoV-2 PLpro catalytic site (PDB ID 6WX4) | 10 |
| 9 | **Figure 9S.** Docked poses 1 and 2 of **16** in the SARS-CoV-2 PLpro catalytic site (PDB ID 6WX4) | 11 |
| 10 | **Figure 10S.** Docked poses 1 and 2 of **17** in the SARS-CoV-2 PLpro catalytic site(PDB ID 6WX4) | 12 |
| 11 | **Figure 11S.** Docked poses 1 and 2 of **19** in the SARS-CoV-2 PLpro catalytic site(PDB ID 6WX4) | 13 |
| 12 | **Figure 12S.** Docked poses 1 and 2 of **20** in the SARS-CoV-2 PLpro catalytic site(PDB ID 6WX4) | 14 |

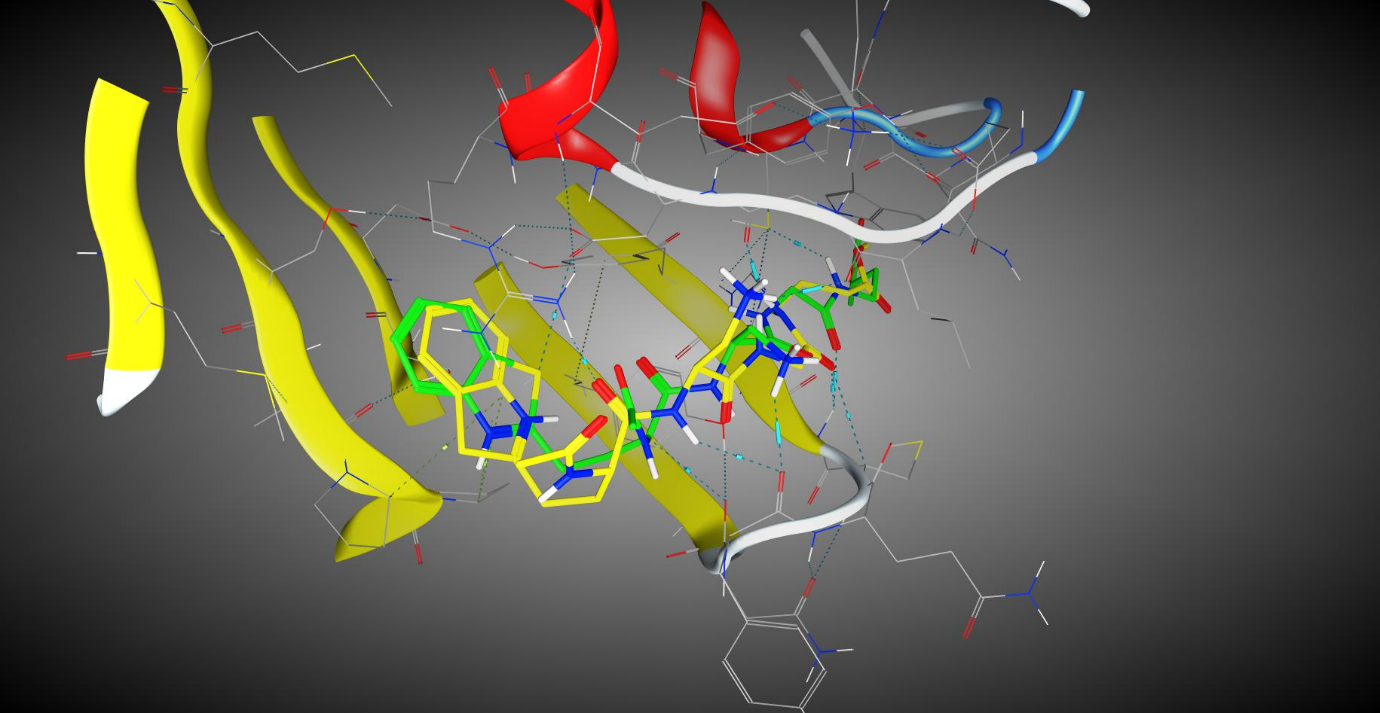
a.

b.	c.

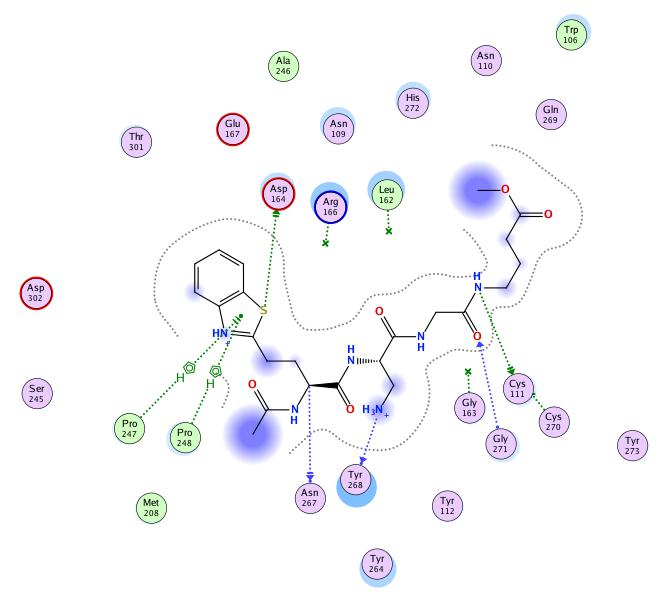

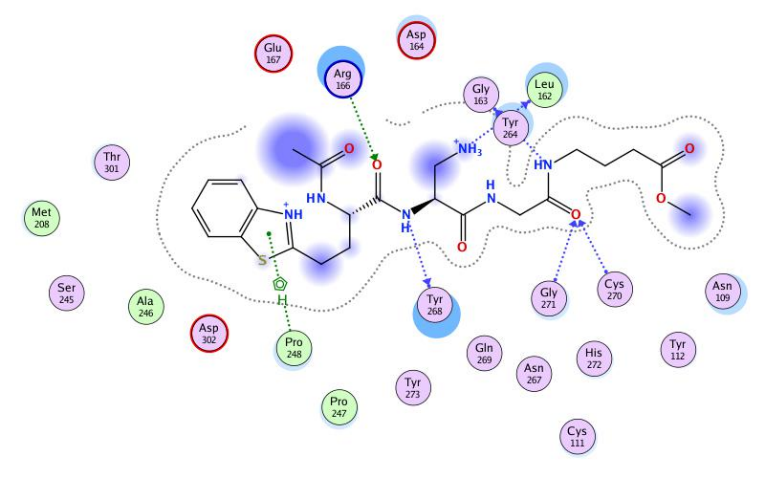

**Figure 1S.** Docked poses of crystal structure ligand (PDB ID 6WUU) (Pose 1 - green capped-stick

representation and (Pose 2 - yellow capped stick Representation) in the SARS-CoV-2

PLpro catalytic site (PDB ID 6WUU); 2D interaction diagram of b. Pose 1 and c.

Pose 2 of crystal structure ligand

a.

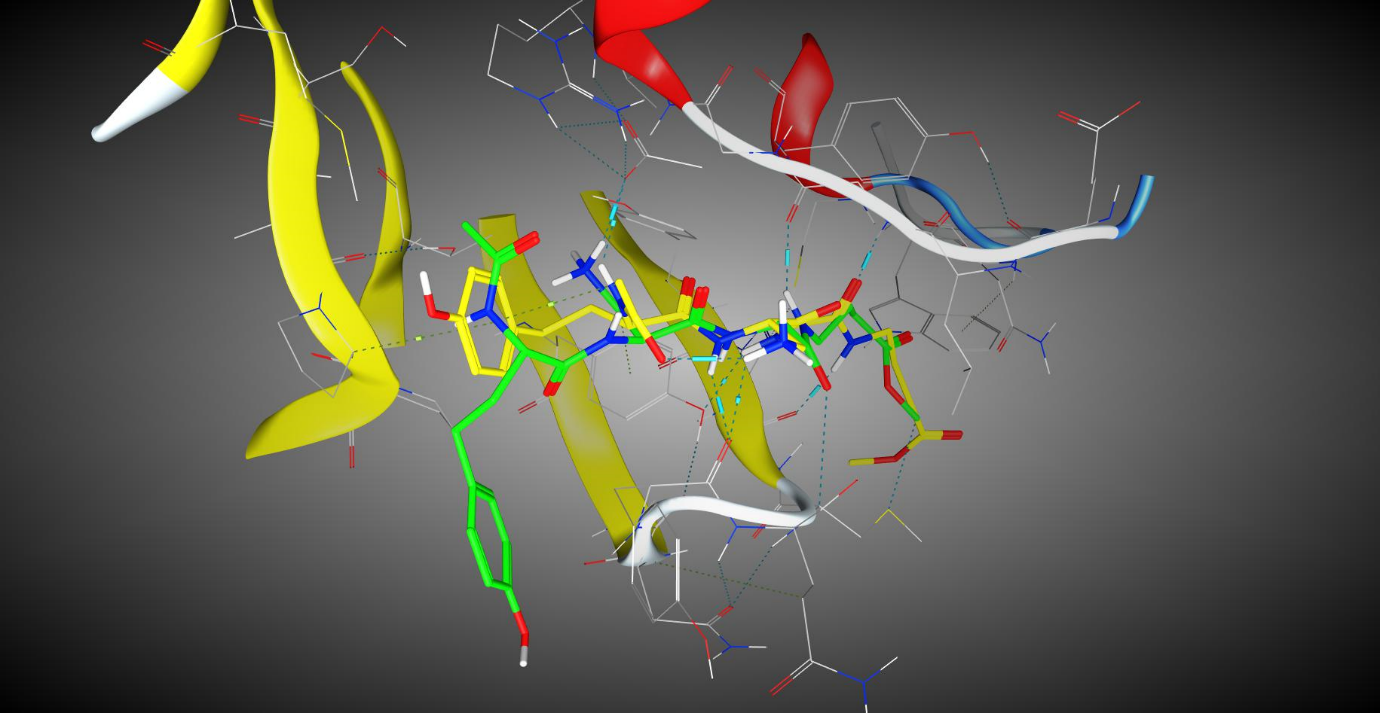

b.	c.

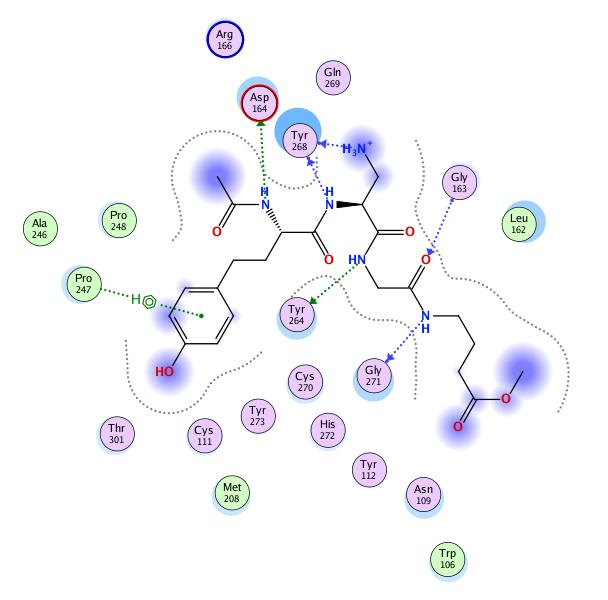

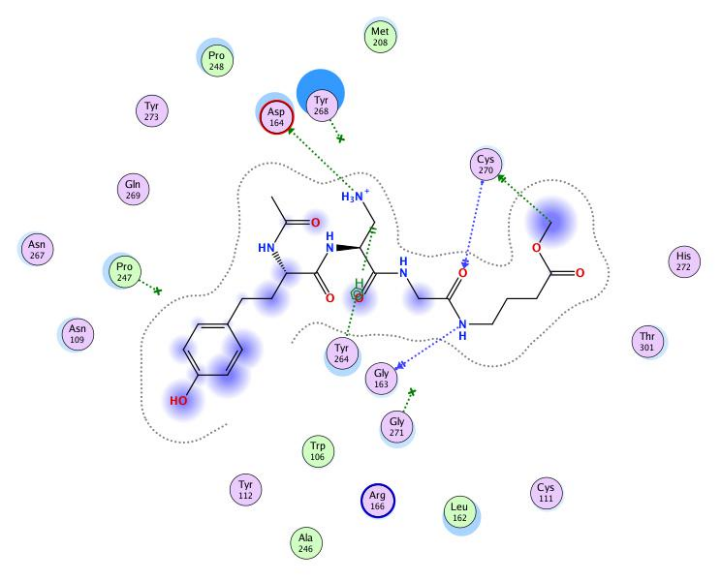

**Figure 2S.** Docked poses of crystal structure ligand (PDB ID 6WX4) (Pose 1 - green capped-stick

representation and (Pose 2 - yellow capped stick Representation) in the SARS-CoV-2

PLpro catalytic site (PDB ID 6WX4); 2D interaction diagram of b. Pose 1 and c.

Pose 2 f crystal structure ligand

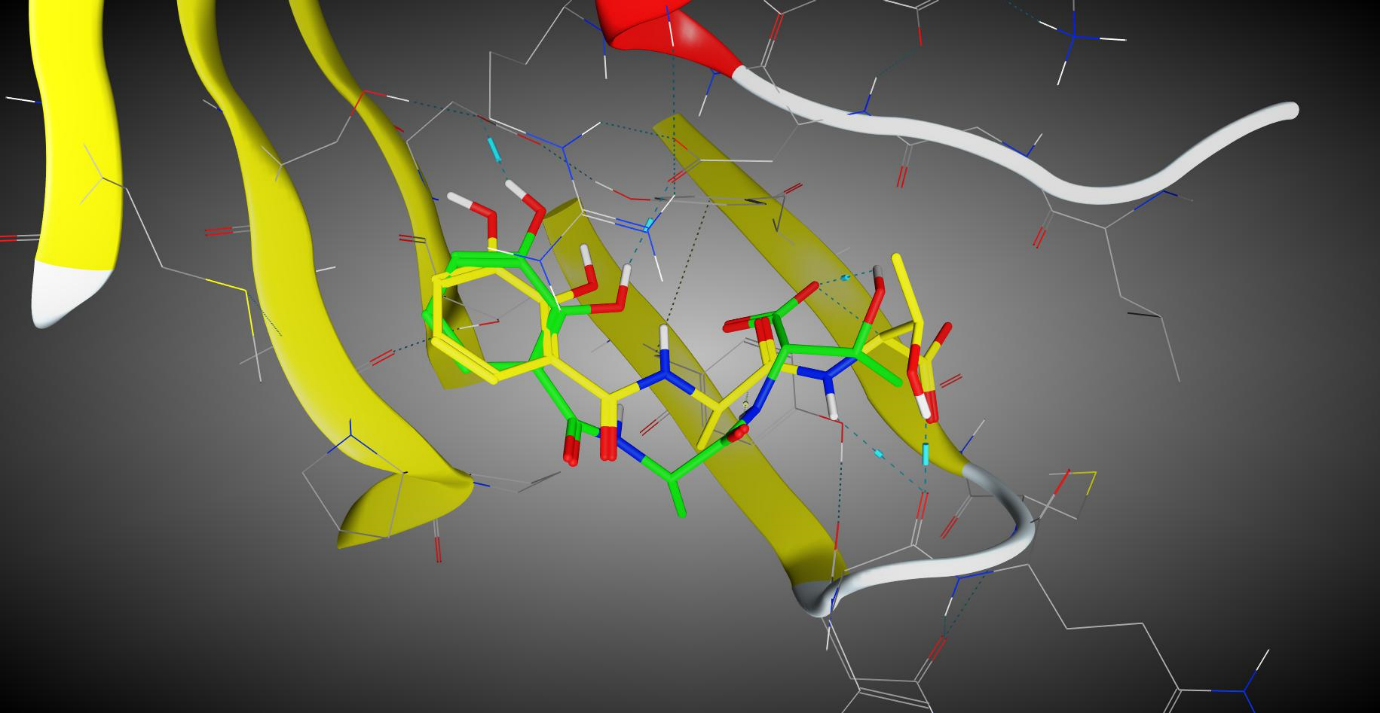
a.

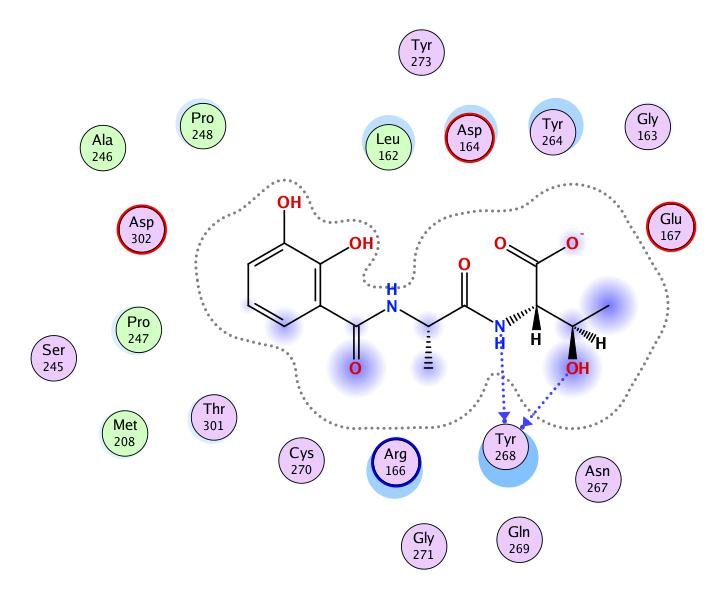

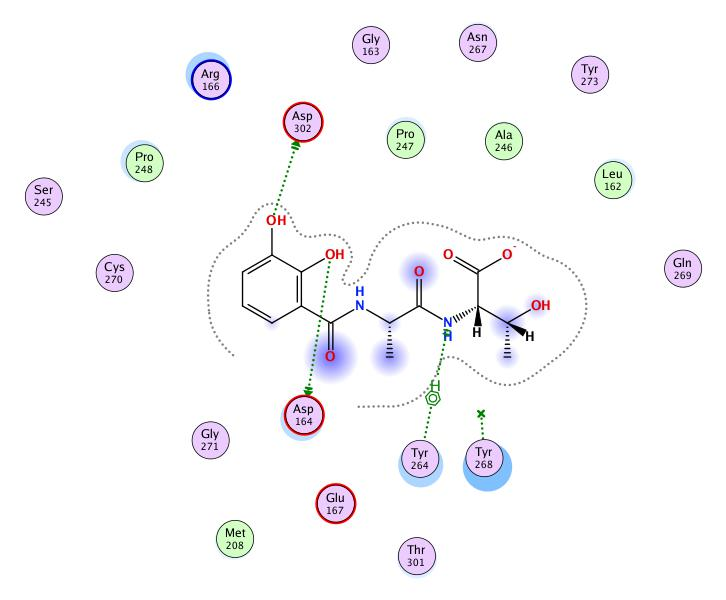
b.	c.

**Figure 3S.** Docked poses 1 (green capped-stick representation) and 2 (yellow capped stick

representation) of **13** in the SARS-CoV-2 PLpro catalytic site (PDB ID 6WUU); 2D

interaction diagram of b. **13** – Pose 1 and c. **13** – Pose 2

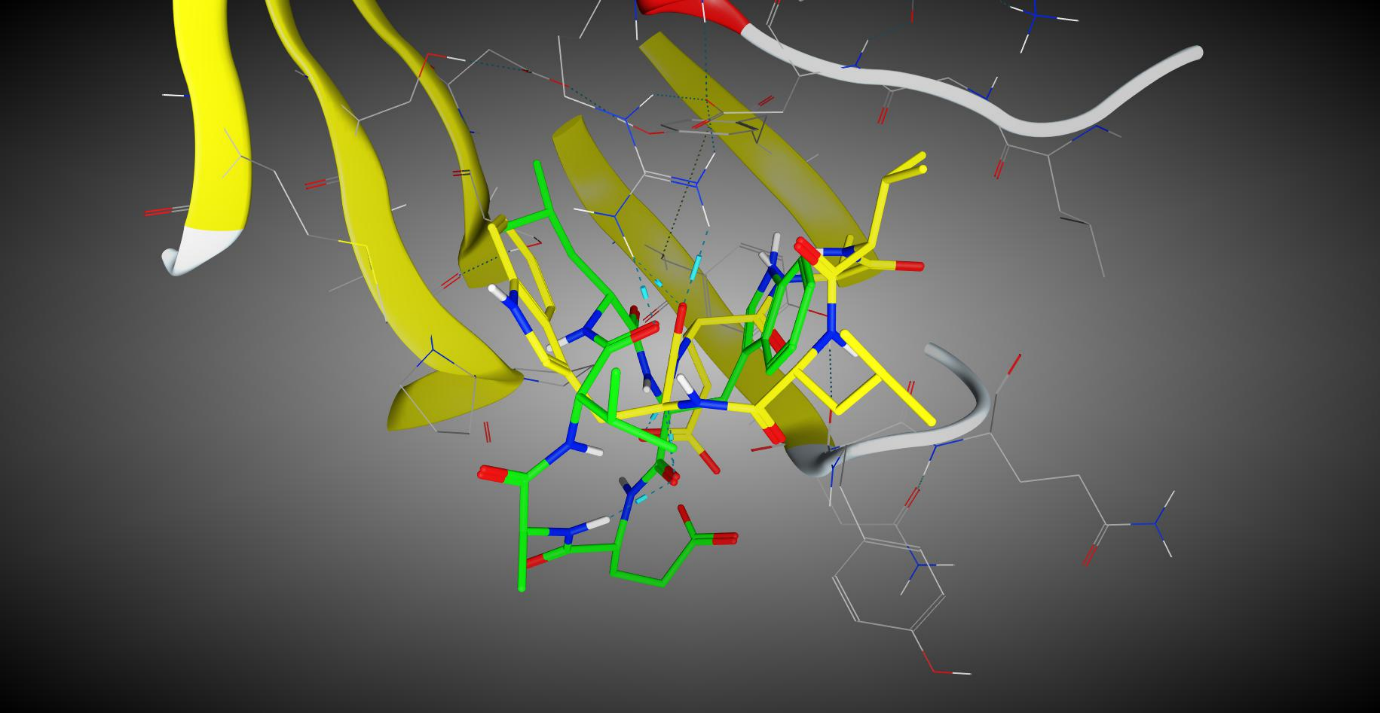
a.

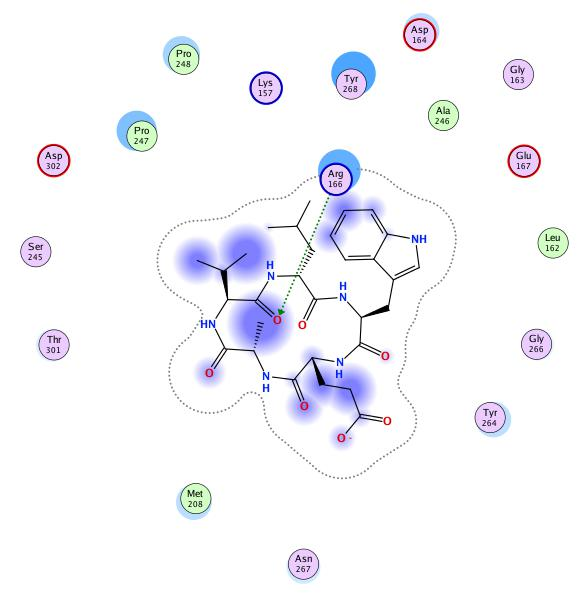
b.	c.

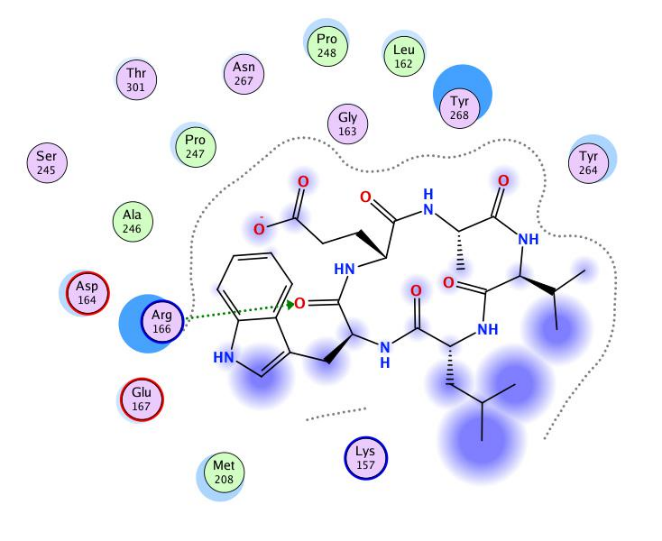

**Figure 4S.** Docked poses 1 (green capped-stick representation) and 2 (yellow capped stick

representation) of **14** in the SARS-CoV-2 PLpro catalytic site (PDB ID 6WUU); 2D

interaction diagram of b. **14** – Pose 1 and c. **14** – Pose 2

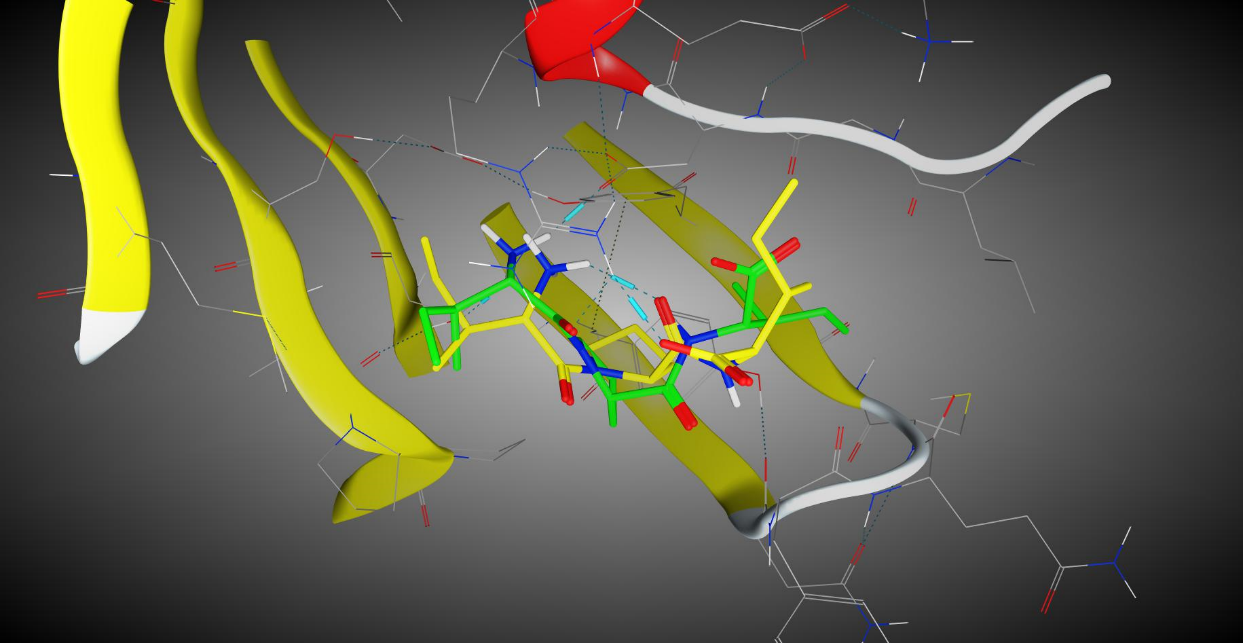
a.

b.	c.

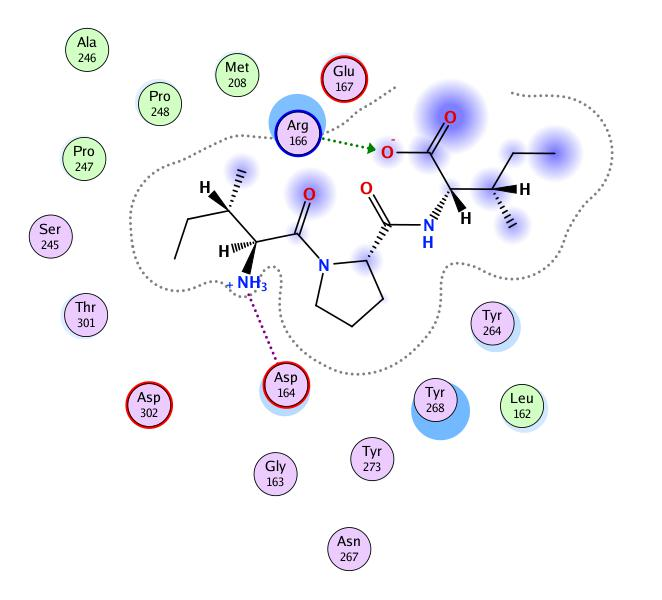

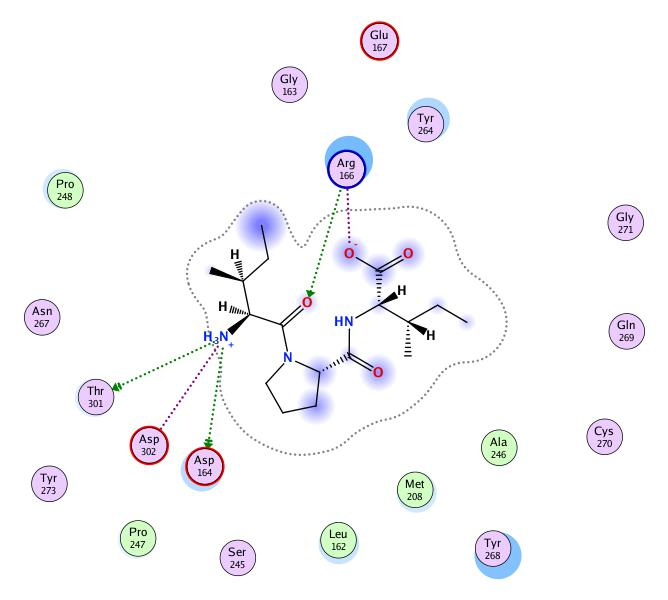

**Figure 5S.** Docked poses 1 (green capped-stick representation) and 2 (yellow capped stick

representation) of **15** in the SARS-CoV-2 PLpro catalytic site (PDB ID 6WUU); 2D

interaction diagram of b. **15** – Pose 1 and c. **15** – Pose 2

a.

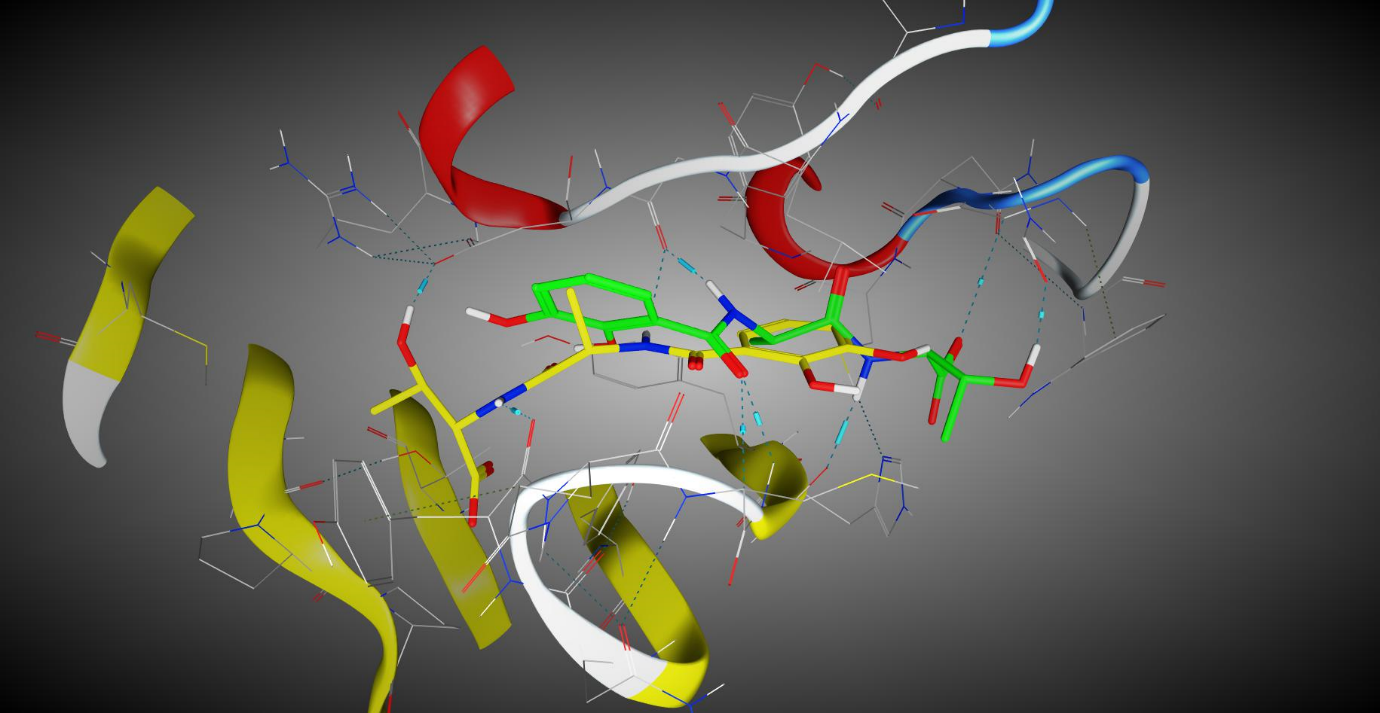

b.	c.

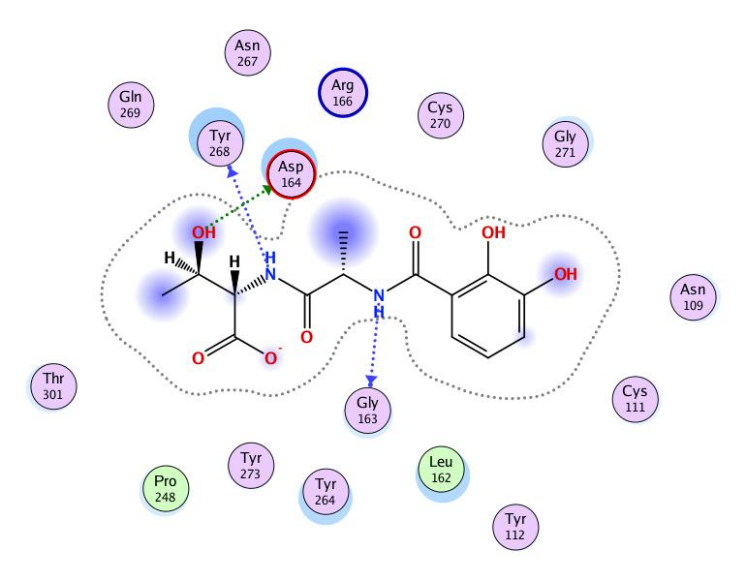

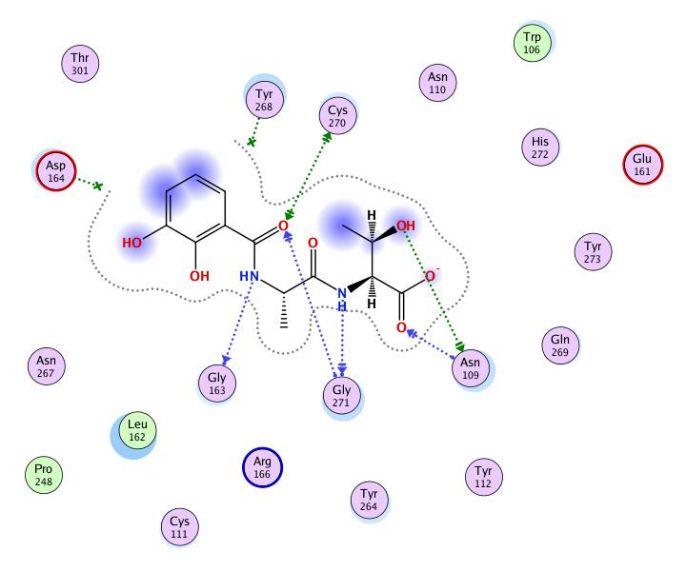

**Figure 6S.** Docked poses 1 (green capped-stick representation) and 2 (yellow capped stick

representation) of **13** in the SARS-CoV-2 PLpro catalytic site (PDB ID 6WX4); 2D

interaction diagram of b. **13** – Pose 1 and c. **13** – Pose 2

a.

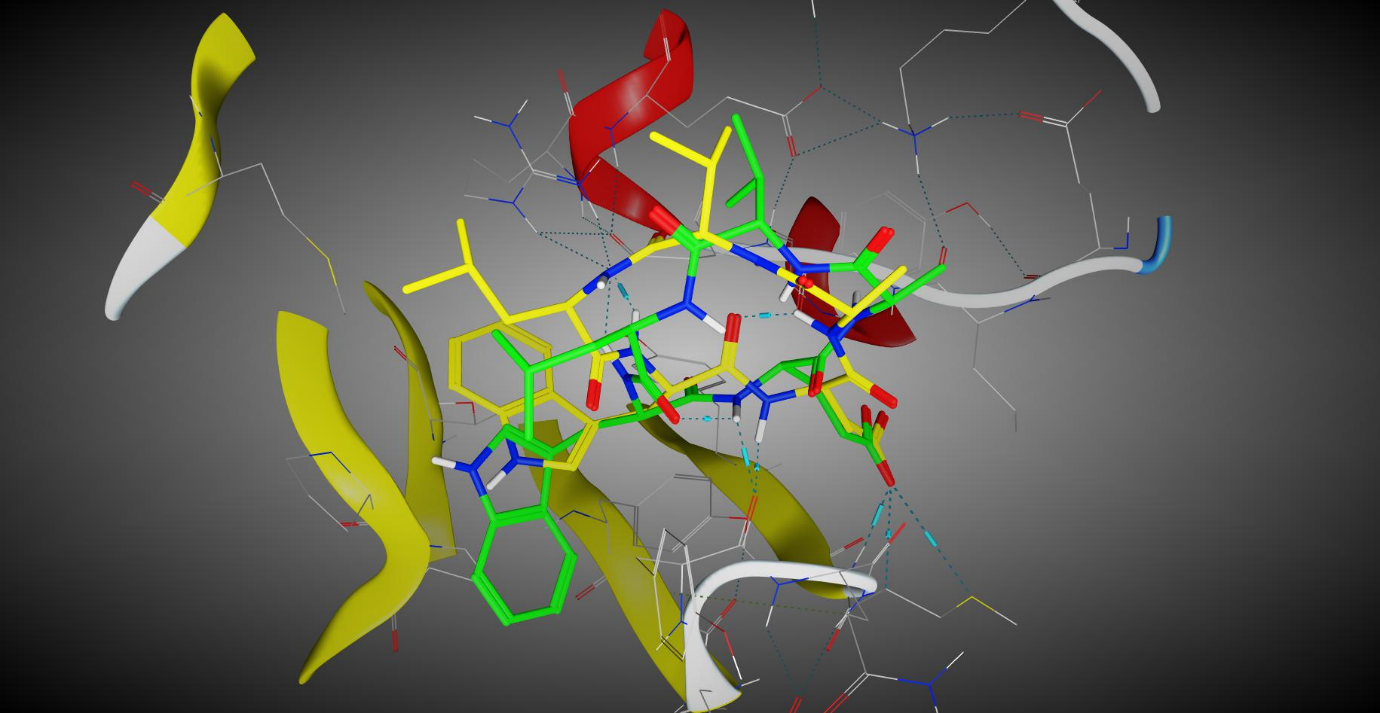

b.	c.

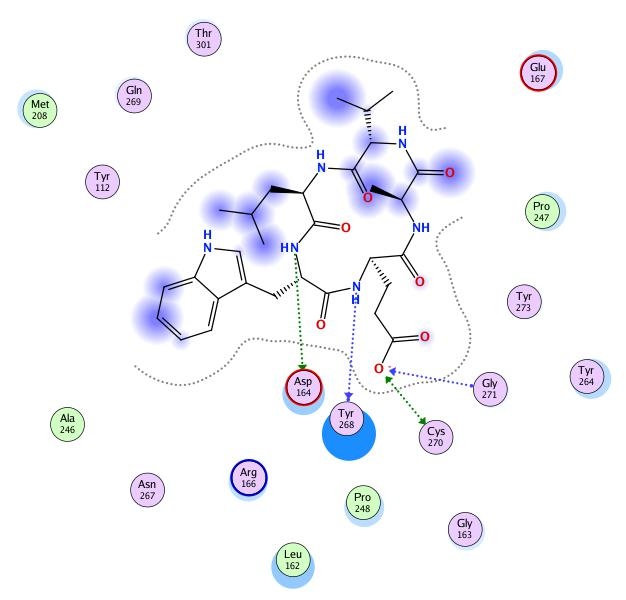

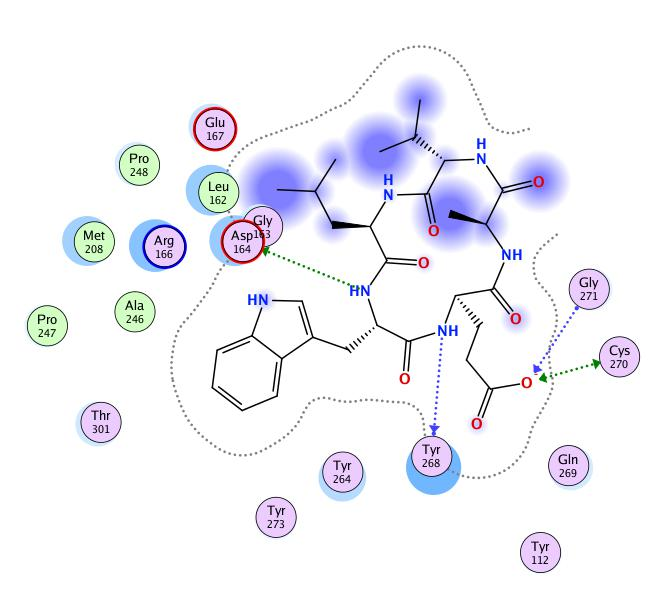

**Figure 7S.** Docked poses 1 (green capped-stick representation) and 2 (yellow capped stick

representation) of **14** in the SARS-CoV-2 PLpro catalytic site (PDB ID 6WX4); 2D

interaction diagram of b. **14** – Pose 1 and c. **14** – Pose 2

a.

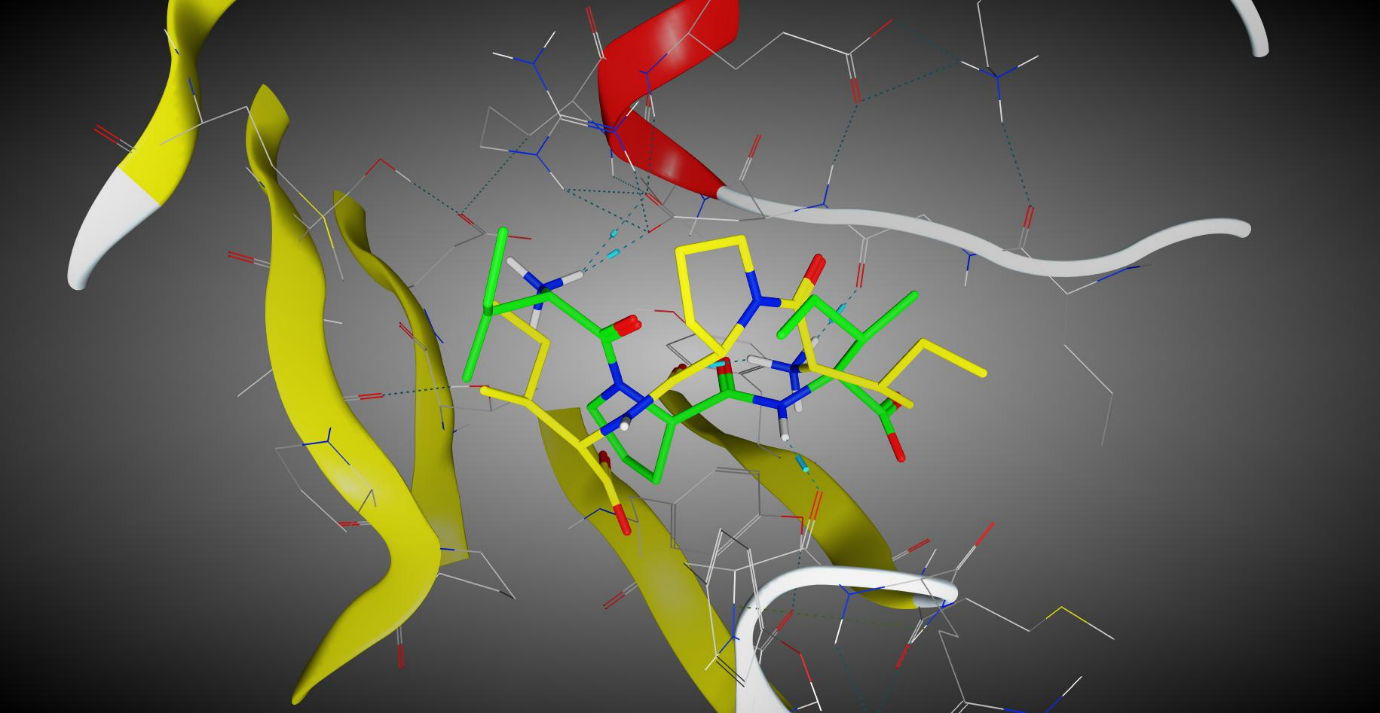

b.	c.

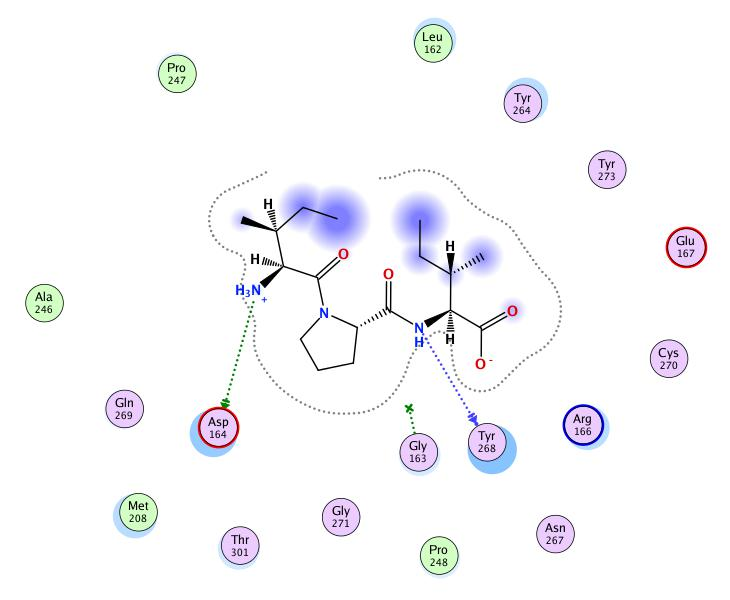

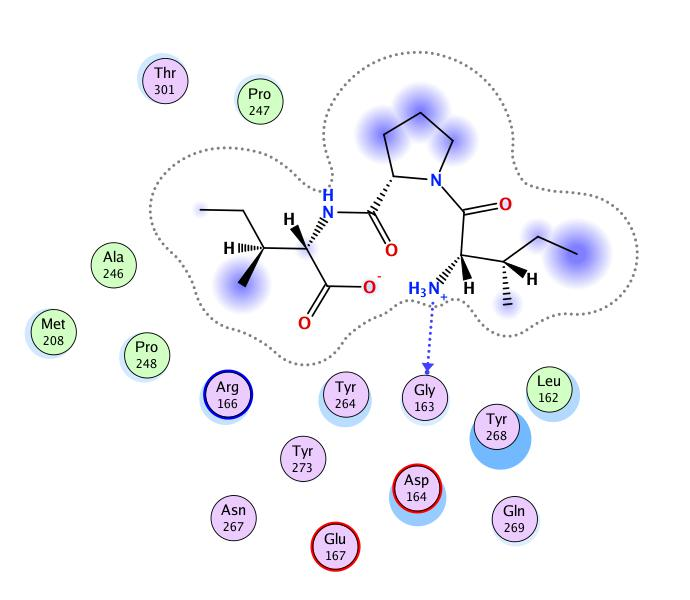

**Figure 8S.** Docked poses 1 (green capped-stick representation) and 2 (yellow capped stick

representation) of **15** in the SARS-CoV-2 PLpro catalytic site (PDB ID 6WX4); 2D

interaction diagram of b. **15** – Pose 1 and c. **15** – Pose 2

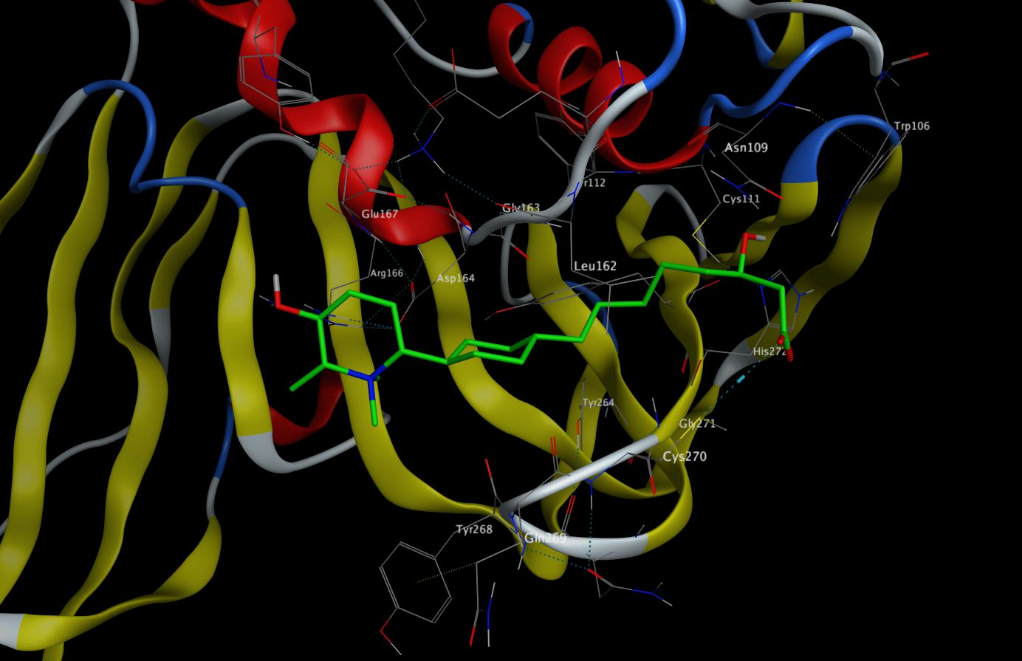
a.

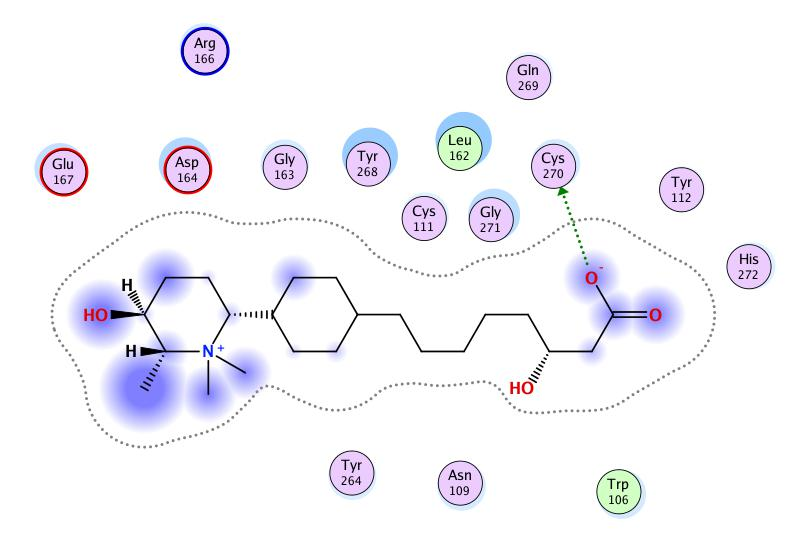
b.

**Figure 9S.** Docked pose 1 (green capped-stick representation) of **16** in the SARS-CoV-2 PLpro

catalytic site (PDB ID 6WX4); 2D interaction diagram of b. **16** – Pose 1 with binding-site residues

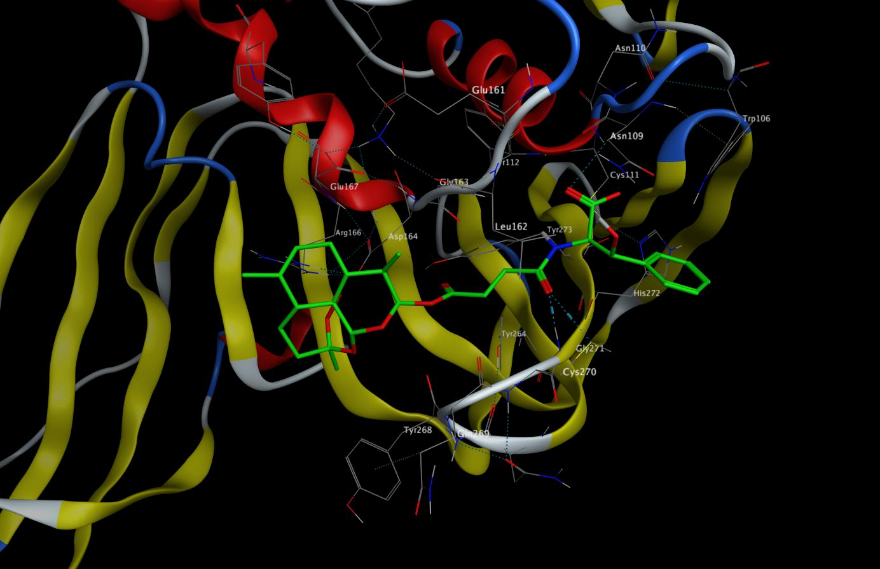
a.

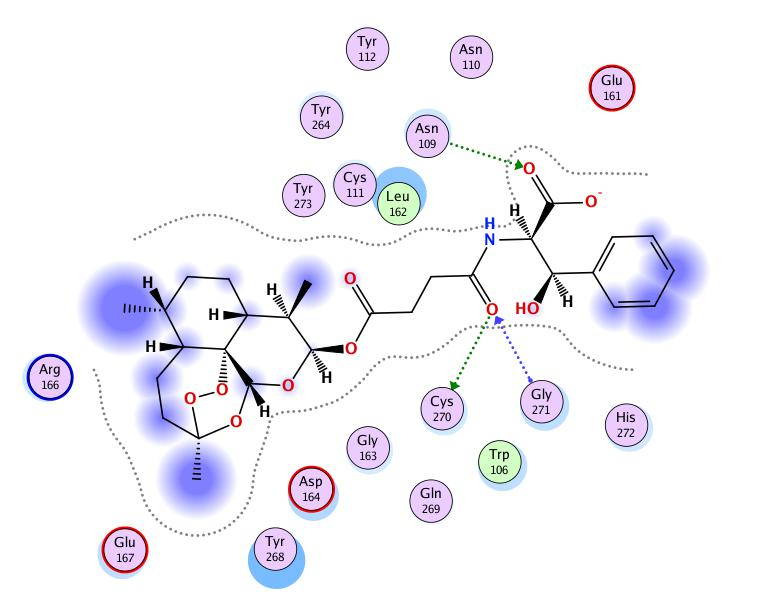
b.

**Figure 10S.** Docked pose 1 (green capped-stick representation) of **17** in the SARS-CoV-2 PLpro

catalytic site (PDB ID 6WX4); 2D interaction diagram of b. **17** – Pose 1 with binding-site

residues

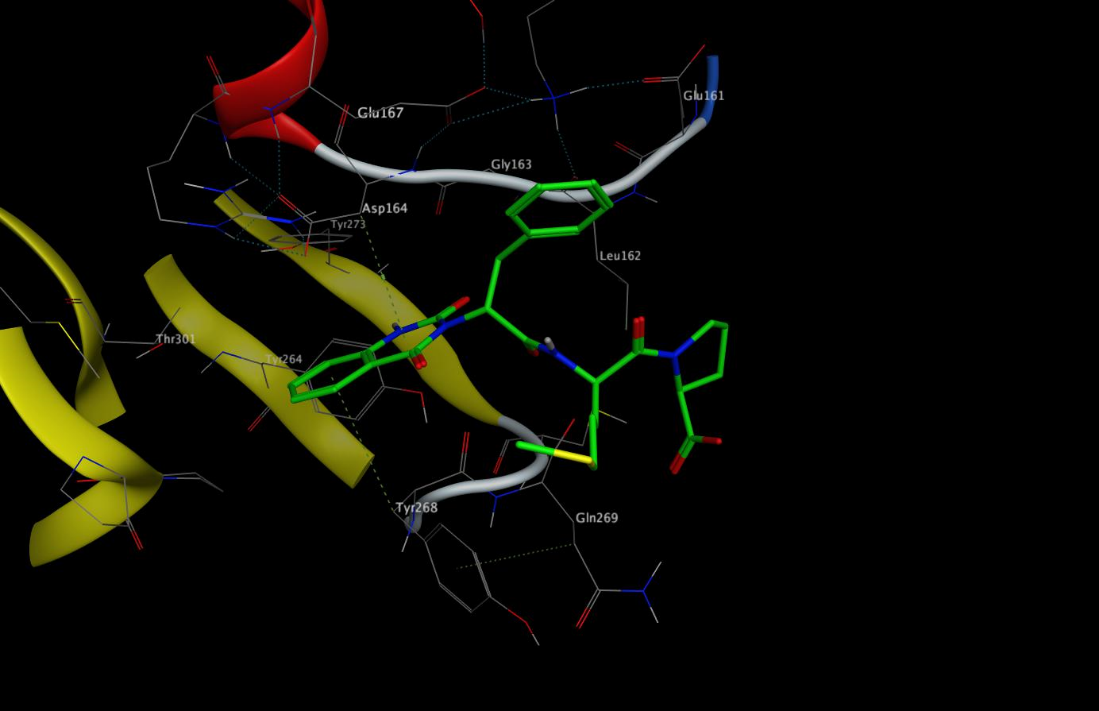
a.

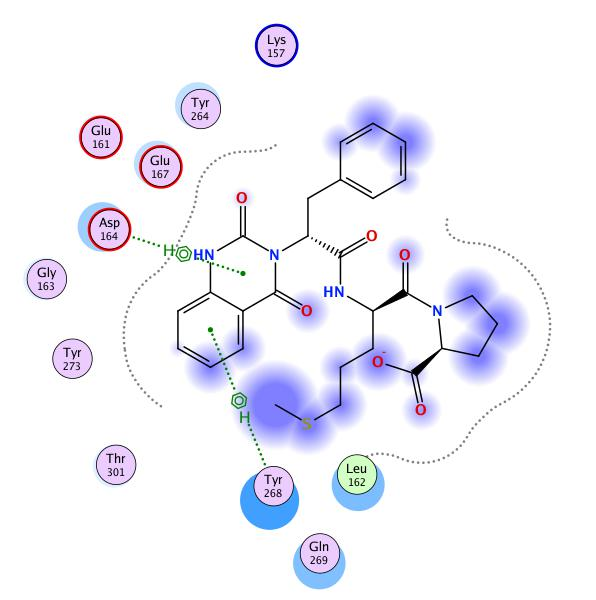
b.

**Figure 11S.** Docked pose 1 (green capped-stick representation) of **19** in the SARS-CoV-2 PLpro

catalytic site (PDB ID 6WX4); 2D interaction diagram of b. **19** – Pose 1 with binding-site

residues

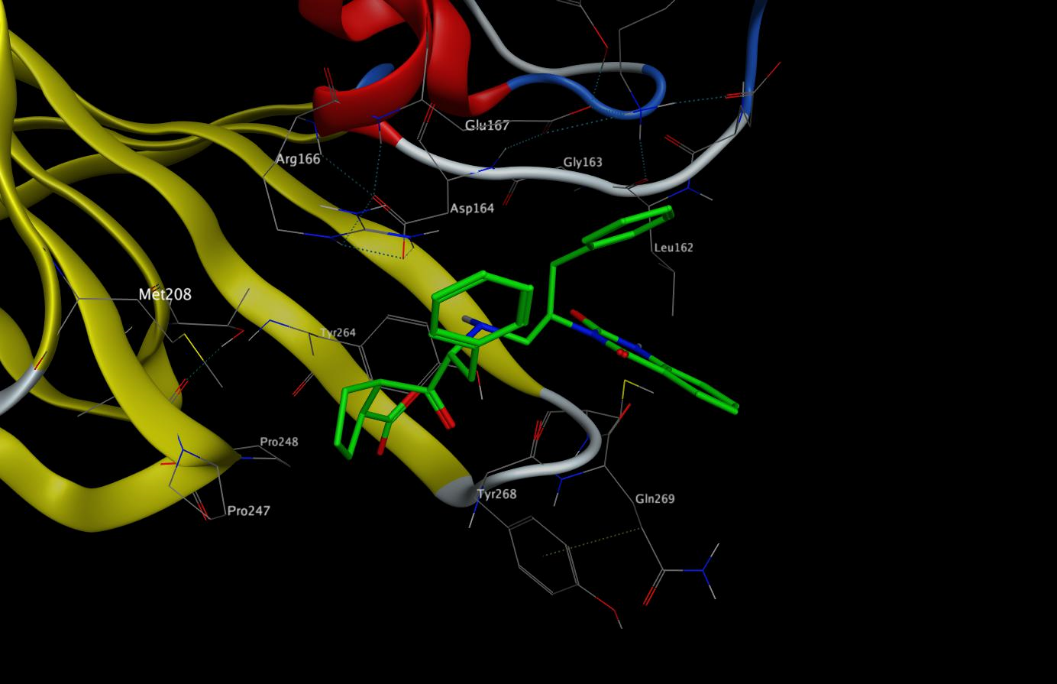
a.

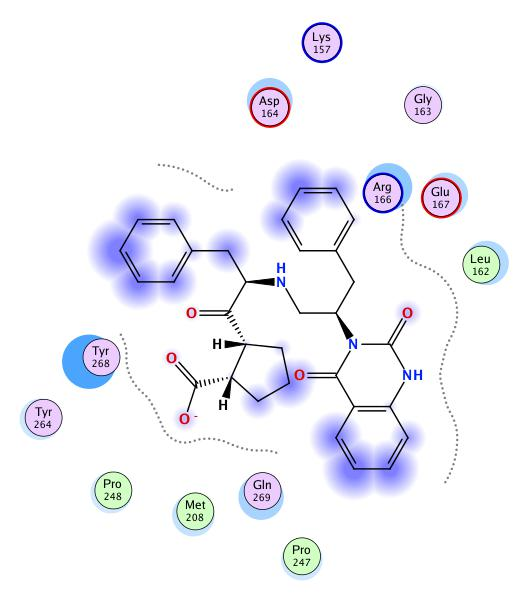
b.

**Figure 12S.** Docked pose 1 (green capped-stick representation) of **20** in the SARS-CoV-2 PLpro

catalytic site (PDB ID 6WX4); 2D interaction diagram of b. **20** – Pose 1 with binding-site

residues
